# Structural mechanism of Necrocide 1 activation of human TRPM4 that triggers necrosis by sodium overload

**DOI:** 10.64898/2026.01.28.702369

**Authors:** Celso M. Teixeira-Duarte, Wan Fu, Weizhong Zeng, Jianghuang Wang, Xinzhe Jiang, Ziye Zhao, Qing Zhong, Youxing Jiang

**Author notes:** These authors contributed equally (Celso M. Teixeira-Duarte, Wan Fu). Correspondence to: Youxing Jiang, Ph.D., Department of Physiology, UT Southwestern Medical Center, 5323 Harry Hines Blvd., Dallas, Texas 75390-9040, Qing Zhong, Ph.D., Department of Pathophysiology, Shanghai Jiao Tong University School of Medicine.

## Abstract

The small molecule Necrocide 1 (NC1) constitutively activates human TRPM4, triggering Na⁺ influx and leading to necrotic cell death, a process termed Necrosis by Sodium Overload (NECSO). NC1 activation is specific to human TRPM4 and does not affect most of the other mammalian TRPM4 orthologs. Here, we elucidate the molecular mechanism underlying NC1 activation and its species-specific selectivity for human TRPM4 using a combination of single-particle cryo-EM, electrophysiology, and cell death assays. We identify the NC1-binding site and the key molecular determinants responsible for channel activation. In addition, we explain the insensitivity of mouse TRPM4 to NC1 and pinpoint specific residues that define NC1 specificity for human TRPM4. Given the upregulation of TRPM4 in various human cancers, our mechanistic insights into NC1 activation and specificity provide a framework for the potential development of cancer therapeutics targeting TRPM4-mediated necrosis.

## Introduction

TRPM4 is a Ca^2+^-activated, monovalent-selective cation channel belonging to the transient receptor potential melastatin (TRPM) subfamily^1–3^. It is ubiquitously expressed^4^ and plays a key role in regulating various physiological processes, including cardiac conduction^5–8^, immune response^9^, and cell death^10–14^. Notably, TRPM4 is upregulated in several cancer types^15^, such as prostate^16–18^, breast^19,20^, diffuse large B-cell lymphoma^21^, and acute myeloid leukemia^22^, making it an attractive target for therapeutic intervention. Despite this potential, commonly used TRPM4 modulators suffer from low specificity^5,23–27^, prompting extensive efforts to develop monoclonal antibodies^28–31^ and small molecules^32,33^ specific for this channel. Among the latter, anthranilic acid derivatives, CBA, NBA, and IBA, have emerged as promising inhibitors^32,34,35^. Recent structural studies revealed that NBA and IBA bind in the vanilloid binding pocket, adjacent to the binding sites of the exogenous modulator decavanadate and the endogenous ligand PI(4,5)P₂, to exert their inhibitory effects^27,35,36^.

TRPM4 is activated by intracellular calcium and PI(4,5)P₂, promoting membrane depolarization via monovalent cation influx^1,23^. Interestingly, TRPM4 channel opening has been implicated in necrotic cell death triggered by anticancer treatments in human breast cancer cells^12^. Necrocide 1 (NC1) ((S’)-3-cycloheptyl-3-(4-hydroxyphenyl)-6-methoxy-7-methyl-1,3-dihydroindole-2-one) is a small molecule derived from the optimization of a compound identified in a screen for anti-tumorigenic activity^37,38^. NC1 induces necrotic cell death in several cancer cell lines, including PC-3 (prostate) and MCF-7 (breast), and has been shown to suppress tumor growth in human cancer xenografts^39^. Importantly, NC1 does not trigger necrosis in non-tumorigenic cells such as fibroblasts and umbilical vein endothelial cells^39^.

Recent findings demonstrate that NC1 induces necrotic cell death by directly activating human TRPM4, triggering Na⁺ influx that leads to osmotic swelling and membrane rupture, a process termed Necrosis by Sodium Overload (NECSO)^40,41^. Interestingly, NC1 exhibits a high degree of species specificity: it activates human TRPM4 but not most other mammalian TRPM4 orthologues or other human TRPM family members. Since evading apoptosis is a hallmark of cancer cells^42,43^, targeting alternative, non-apoptotic pathways such as TRPM4-mediated necrosis may offer novel therapeutic avenues. Understanding the molecular mechanism of NC1-induced human TRPM4 activation and its species specificity is therefore critical for the rational development of TRPM4-targeted anticancer therapies.

In this study, we combine single-particle cryo-EM, electrophysiology, and cell death assays to elucidate the structural mechanism by which NC1 activates human TRPM4. We demonstrate that NC1 functions as a non-competitive surrogate of the endogenous Ca^2+^ ligand – it binds to a pocket within the S1-S4 domain adjacent to the Ca^2+^ site and induces the same conformational changes as those triggered by Ca^2+^. Like Ca^2+^-mediated activation, NC1-induced channel opening also requires membrane PI(4,5)P_2_ to stabilize the open state. Through comparative mutagenesis and structural analysis of human and mouse TRPM4, we identify the molecular determinants of NC1 specificity. Our results reveal that the insensitivity of mouse TRPM4 to NC1 arises not from a lack of binding, but from drug-induced conformational changes that destabilize the selectivity filter and inactivate the channel. We identify three critical residues that confer NC1 sensitivity, and their substitution renders mouse TRPM4 responsive to NC1, offering insights into the species selectivity of NC1 and paving the way for future therapeutic development.

## Results

### NC1 activates hTRPM4 and promotes cell death

The small molecule Necrocide 1 (NC1) was recently shown to induce necrotic cell death by activating the human TRPM4 (hTRPM4) channel^40^. The key electrophysiological features of NC1-activated hTRPM4 were characterized using whole-cell patch-clamp recordings of hTRPM4-expressing HEK293 cells (Methods). NC1 activates TRPM4 slowly, requiring about two minutes for the current to reach its maximum, indicating a slow on-rate of NC1 binding (Fig. 1a and 1b). Most channels remain activated even after several minutes of washout, suggesting high-affinity binding and an extremely slow off-rate. With the presence of 1 mM EGTA in the pipette solution (cytosolic side), our recording conditions eliminate the potential contribution to channel activation from cytosolic Ca²⁺, the endogenous ligand of TRPM4, confirming the Ca²⁺-independence of NC1 activation.

**Fig. 1:**
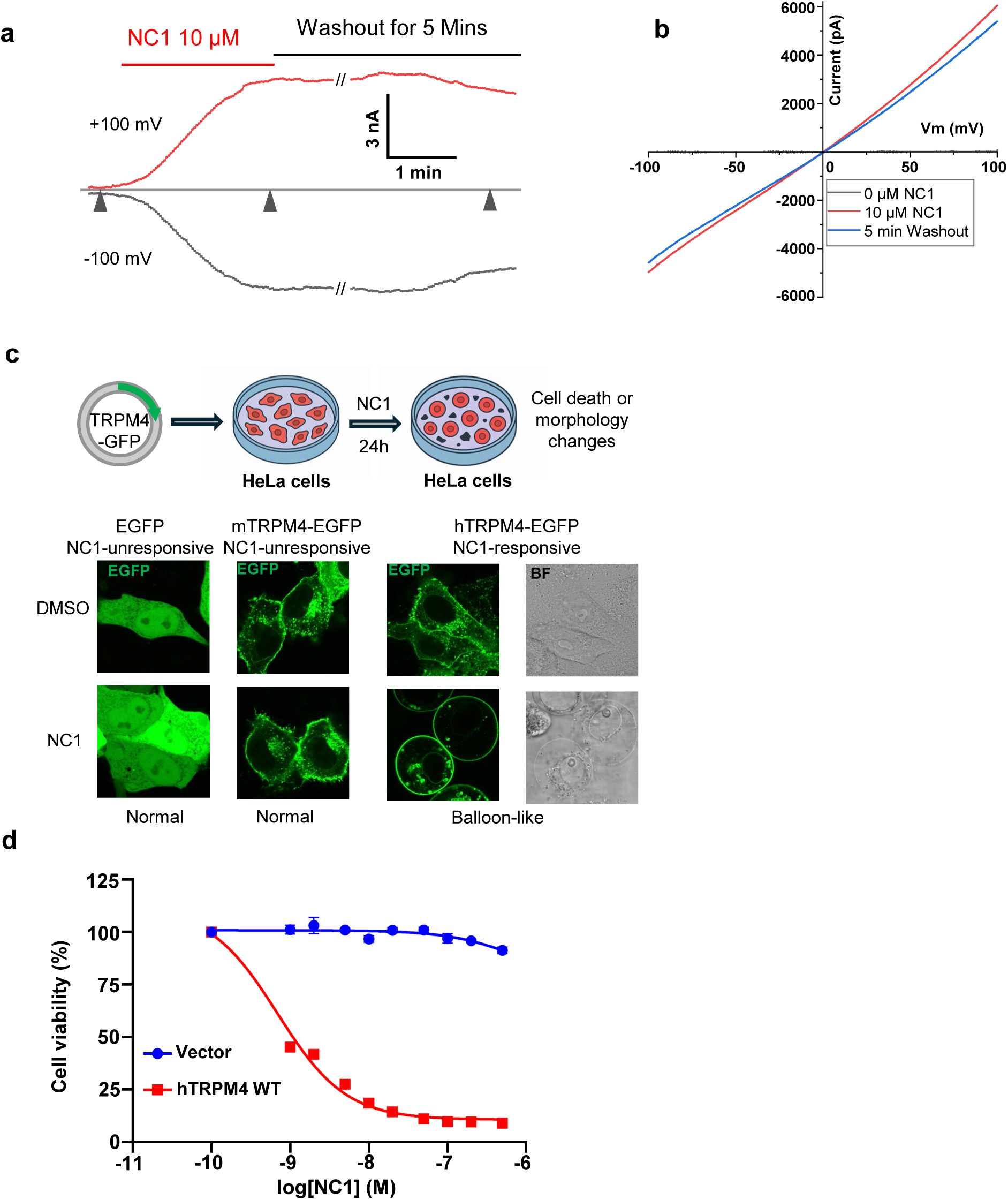
NC1 activation of hTRPM4. **a** Macroscopic currents of hTRPM4 overexpressed in HEK293 cells at ±100 mV in a whole-cell patch with the presence or absence of 10 µM NC1 in the bath (extracellular). Triangles mark the time points of three NC1-mediated hTRPM4 activation states (left to right): Closed channel in the absence of NC1; Fully activated channel 2 min after NC1 addition; Channel current retained 5 min after NC1 washout. **b** Sample I-V curves measured at the time points marked by the triangles shown in **a**. **c** NC1-induced cell death assay. Top panel: schematic illustration of the assay. Bottom panels: morphologies of EGFP, EGFP-tagged mTRPM4, or EGFP-tagged hTRPM4-expressing HeLa cells 24 hours after DMSO (NC1 solvent) or NC1 (1 µM) treatment. **d** Dose-dependent NC1 cytotoxicity in HeLa cells stably expressing wild-type hTRPM4. Data points represent mean ± SD of n = 3 independent replicates.

To complement electrophysiological data, we conducted cell death and viability assays to assess hTRPM4-dependent necrosis in HeLa cells (Methods). While HeLa cells are insensitive to NC1 due to a lack of detectable endogenous TRPM4, they become susceptible to NC1-induced necrosis upon transient or stable expression of hTRPM4. In the cell death assay, EGFP-tagged hTRPM4 was transiently expressed in HeLa cells, and necrotic morphology, characterized by ‘balloon-like’ cell swelling, was observed 24 hours after NC1 treatment (Fig. 1c). No necrosis was observed in the NC1-treated control cells transiently expressing EGFP or EGFP-tagged mouse TRPM4. This assay provides a qualitative readout of hTRPM4-dependent necrosis, enabling the screening of mutational effects on the NC1 sensitivity of the channel, as discussed later. In the cell viability assay, hTRPM4 was stably expressed in HeLa cells. Following 24-hour NC1 treatment at varying concentrations, cytotoxicity was assessed by measuring intracellular ATP levels, indicative of cell viability, using the CellTiter-Glo (CTG) assay (Fig. 1d, Methods). This assay provides a quantitative measure of NC1’s dose-dependent toxicity in hTRPM4-expressing cells.

### Structure of NC1-bound TRPM4

To elucidate the structural basis of NC1 binding and its activation mechanism, we purified detergent-solubilized hTRPM4 and determined its structure in complex with NC1 at 2.7 Å resolution using single-particle cryo-EM (Fig. 2a, Supplementary Fig. 1a, Supplementary Table 1, and Methods). The NC1 ligand is unambiguously resolved in the EM density map (Fig. 2b). Occupying a pocket within the S1–S4 domain just above the Ca²⁺ activation site, NC1 forms extensive hydrophobic and hydrogen-bonding interactions with surrounding residues (Fig. 2b).

**Fig. 2:**
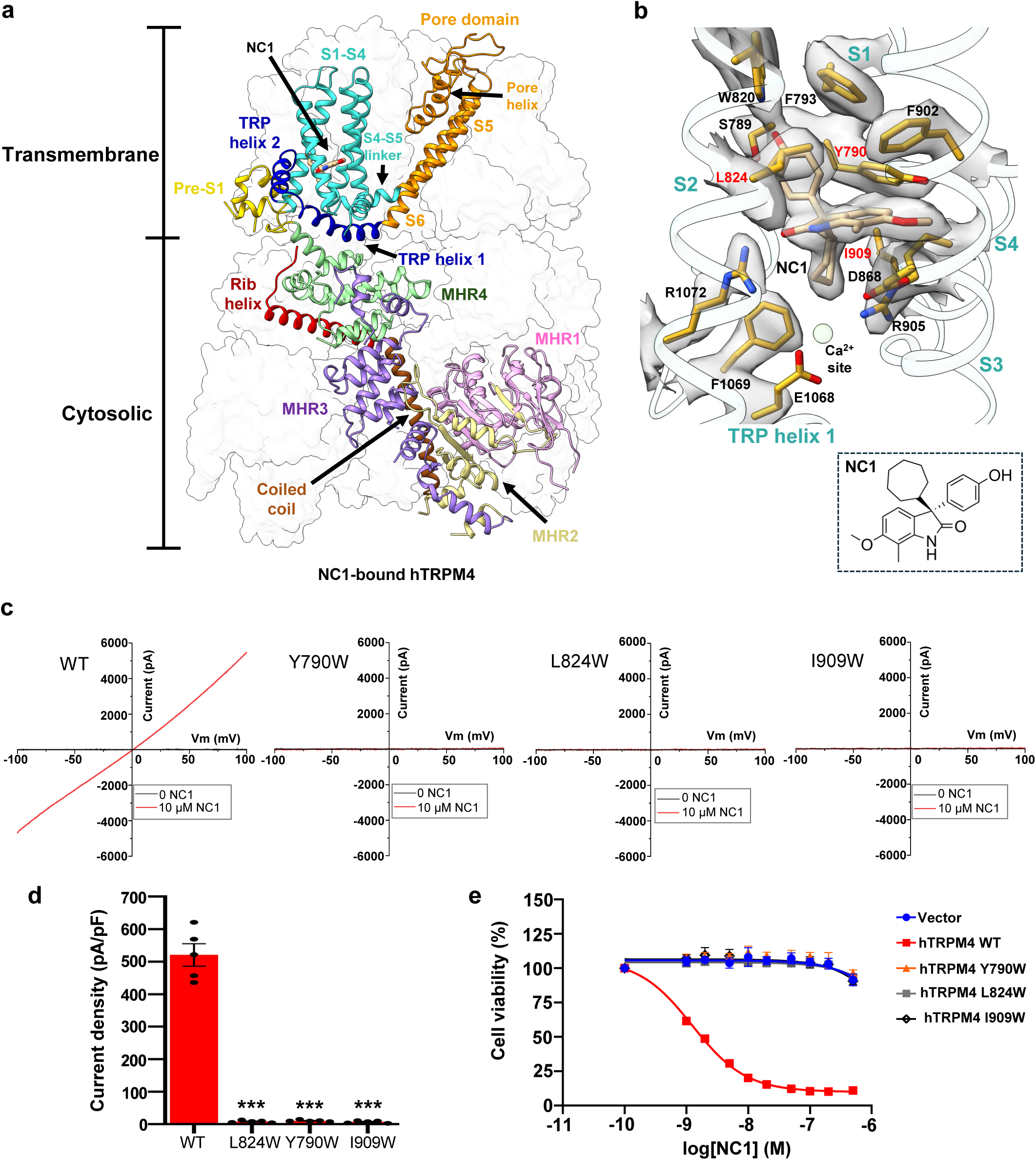
NC1 binding in hTRPM4. **a** Structure of NC1-bound hTRPM4. The front subunit is depicted in a cartoon representation, with each domain individually colored. **b** Zoomed-in view of the NC1 binding site. NC1 and its surrounding residues are shown in stick representation. The light green sphere marks the calcium activation site. Density for NC1 and its surrounding residues (grey surface) is contoured at 0.2 in ChimeraX. Residues sensitive to tryptophan substitution are labeled in red. **c** Sample I-V curves of wild-type hTRPM4 and its tryptophan substitutions at the NC1-binding site recorded in whole-cell patches with 10 µM NC1 in the bath (extracellular). **d** Outward current density of NC1-activated wild-type hTRPM4 and its tryptophan mutations recorded at +100 mV in whole-cell patches with 10 µM NC1 in the bath (extracellular). Bars represent mean ± SEM of n=5 independent replicates (shown as dots). p-values were calculated using a two-sided Student’s t-test and are provided in the Source Data file. *** represents p < 0.001. **e** Dose-dependent NC1 cytotoxicity in HeLa cells stably expressing wild-type hTRPM4 or its tryptophan mutations. Data points represent mean ± SD of n = 3 independent replicates.

To validate NC1 binding, we performed mutagenesis of residues interacting with the ligand. Due to the extensive contact surface between NC1 and its binding pocket, single-alanine substitutions were insufficient to disrupt NC1 binding and channel activation (Supplementary Fig. 2a). We therefore introduced bulkier tryptophan substitutions to block NC1 access sterically. Among all tested, three hTRPM4 single-tryptophan mutants - Y790W, L824W, and I909W – completely lost sensitivity to NC1 (Fig. 2c and 2d). These mutants remained responsive to Ca²⁺/PI(4,5)P_2_ activation, particularly I909W (Supplementary Fig. 2b), confirming that these mutations prevent NC1 binding while retaining native activation. The loss of NC1 activation in these mutants was further supported by cell viability assays using stable HeLa cell lines expressing either wild-type or mutant hTRPM4. While NC1 treatment induced a dose-dependent loss of viability in WT-expressing cells, the mutant-expressing cells were resistant to NC1-induced necrosis (Fig. 2e).

Despite robust NC1 activation observed in functional assays, the NC1-bound TRPM4 structure adopts a conformation intermediate between the closed and open states, when compared to previously determined apo (closed) and Ca²⁺/PI(4,5)P_2_-bound (open) structures (Fig. 3a and Supplementary Fig. 3a). NC1 binding induces local conformational changes in the S1-S4 domain resembling those triggered by Ca²⁺ (Fig. 3b and Supplementary Fig. 3b); however, the channel pore and the cytosolic domains remain in a closed conformation (Fig. 3a, 3c, 3d, and Supplementary Fig. 3a). Within S1-S4, NC1 binding causes the S3 helix to shift inward toward the NC1-binding pocket, moving the W864 side chain closer to the C-terminal end of S4 (Fig. 3b). Through tight packing with H908 on S4, W864 motion displaces the C-terminal part of S4 toward S5 in the adjacent pore domain (Fig. 3b). In the apo TRPM4 structure, F910 at the C-terminal part of S4 forms a key contact with F935 near the cytosolic end of S5 - a coupling essential for transducing Ca²⁺-induced movements into pore opening, as shown in the Ca²⁺/PI(4,5)P_2_-bound structure^36^ (Fig. 3e). In contrast, the NC1-bound structure shows disordered side chains for both F910 and F935, indicating a loss of this critical contact (Fig. 3e). These structural features of NC1-bound hTRPM4, including an activated S1-S4 domain, a closed pore, and disrupted F910–F935 coupling, mirror the Ca²⁺-bound hTRPM4 structure in the absence of PI(4,5)P_2_, previously proposed to represent a desensitized state^36^ (Fig 3e and Supplementary Fig. 3c). Since PI(4,5)P_2_ binding is required to stabilize the open state upon Ca²⁺ activation^36^, the resemblance between NC1- and Ca²⁺-bound structures suggests that NC1 triggers local conformational changes similar to those induced by Ca²⁺ and also relies on PI(4,5)P_2_ binding to stabilize the open channel.

**Fig. 3:**
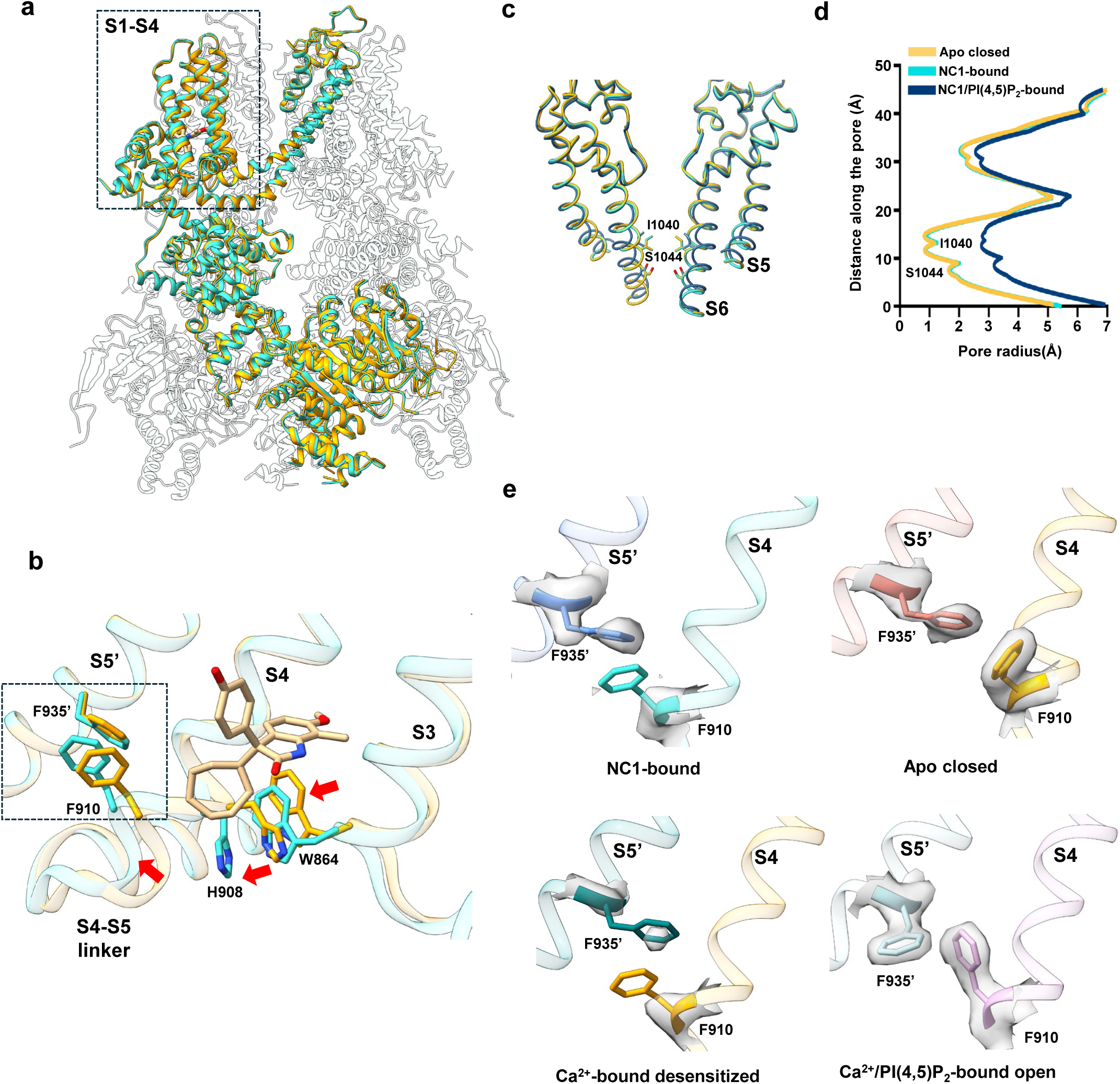
hTRPM4 conformational changes induced by NC1 binding. **a** Superposition of the hTRPM4 structures in the NC1-bound and apo closed (PDB:9MTA) states with the front subunits highlighted in cyan and orange, respectively. **b** Local conformational changes at the S1-S4 domain between NC1-bound (cyan) and apo closed (orange) hTRPM4. Only S3-S4 and the neighboring S5 (marked with a single quotation mark) are displayed for clarity. Red arrows mark the major movements upon NC1 binding. Key residues are shown in stick representation. **c** Structural comparison of the hTRPM4 ion conduction pore in the apo closed (orange), NC1-bound (cyan), and NC1/PI(4,5)P_2_-bound (navy blue) states. Gating residues I1040 and S1044 are shown in stick representation. The front and back subunits are removed for clarity. **d** Pore radius of hTRPM4 along the central axis in the apo closed, NC1-bound, and NC1/PI(4,5)P_2_-bound structures. **e** F935’-F910 contact interface in the NC1-bound, apo closed, Ca^2+^-bound desensitized (PDB:9MT8), and Ca^2+^/PI(4,5)P_2_-bound open (PDB:9MRT) states. The EM density maps for F910 and F935’ are shown in grey surface contoured at 0.23 in ChimeraX.

### PI(4,5)P_2_-dependent NC1 activation of hTRPM4

To test whether PI(4,5)P_2_ is indeed required for NC1 activation, we performed patch-clamp recordings of hTRPM4-expressing HEK293 cells in the inside-out configuration. Following initial activation by cytosolic Ca²⁺, TRPM4 undergoes desensitization to a steady-state level with reduced open probability due to depletion of membrane PI(4,5)P₂, caused by Ca²⁺-induced phospholipase C activation^44,45^ (Fig. 4a). This loss of PI(4,5)P_2_ after Ca²⁺ activation creates a condition suitable for assessing NC1 activation in the absence of PI(4,5)P_2_. Indeed, hTRPM4 becomes unresponsive to NC1 following PI(4,5)P_2_ depletion (Fig. 4a). While PI(4,5)P₂ alone cannot activate TRPM4, its addition in the presence of NC1 robustly activates the channel, restoring current to the pre-desensitization level (Fig. 4a). Consistent with whole-cell recordings, NC1 activates TRPM4 slowly in the presence of PI(4,5)P₂, and the ligand remains tightly bound, making it difficult to wash out (Fig. 4b). These results indicate that PI(4,5)P₂ is a prerequisite for NC1-induced activation of hTRPM4.

**Fig. 4.**
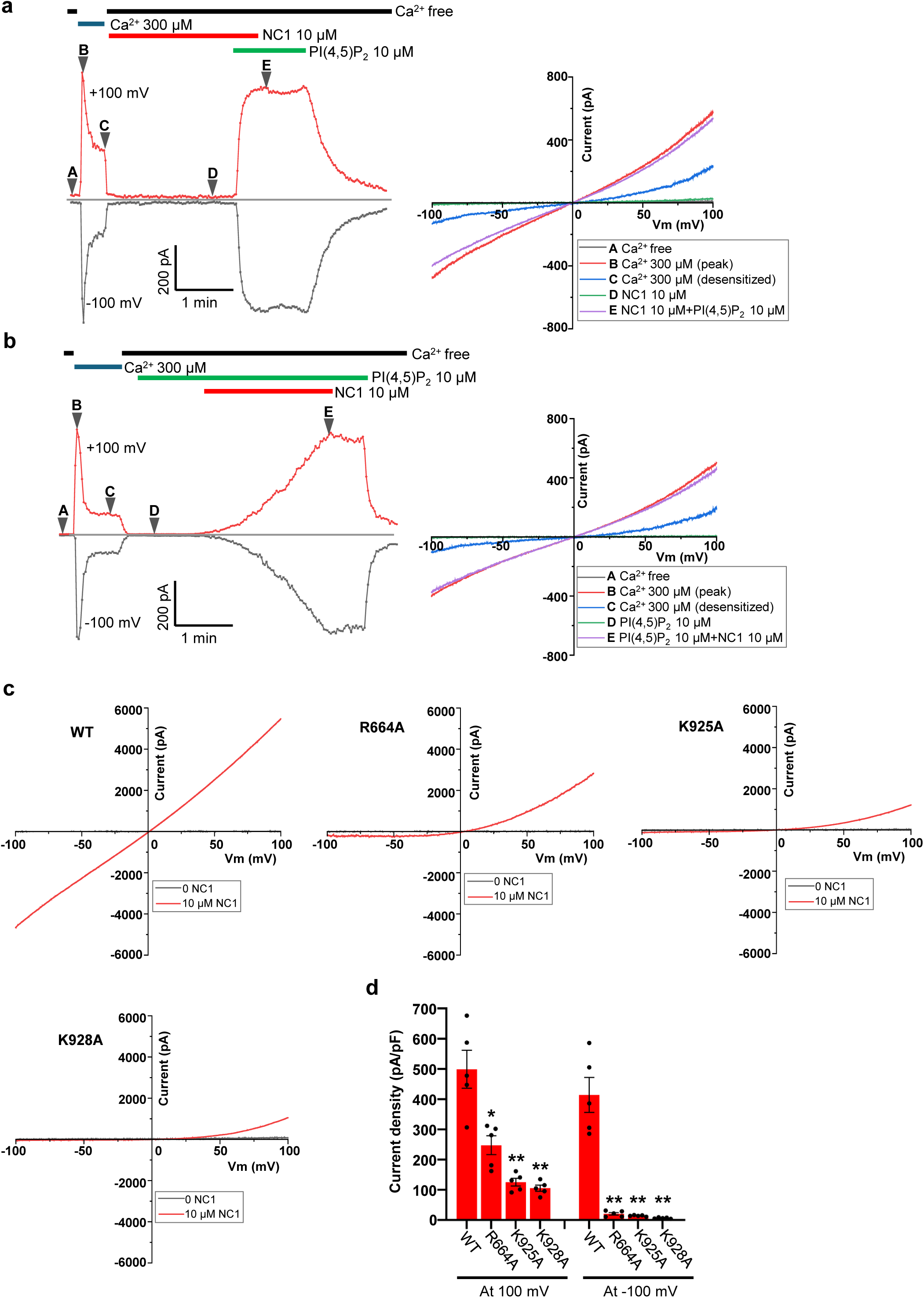
PI(4,5)P_2_-dependent NC1 activation of hTRPM4. **a** Macroscopic currents of hTRPM4-expressing HEK293 cells at ±100 mV in an inside-out patch with the presence or absence of Ca^2+^, NC1, and PI(4,5)P_2_ diC8 in the bath (cytosolic). NC1 and PI(4,5)P_2_ diC8 were introduced after Ca^2+^ activation and desensitization due to Ca^2+^-induced membrane PI(4,5)P_2_ depletion. A-E mark the time points of hTRPM4 in various states: A, apo closed state; B, initial Ca^2+^-activated state before desensitization; C, Ca^2+^-bound desensitized state due to PI(4,5)P_2_ depletion; D, closed state in the presence of NC1; E, NC1/PI(4,5)P_2_-activated state. The panel on the right depicts sample I-V curves corresponding to A-E time points. **b** Similar recordings as **a** except that PI(4,5)P_2_ diC8 was introduced before NC1, and the time point D marks the closed state in the presence of PI(4,5)P_2_ diC8. The panel on the right depicts sample I-V curves corresponding to A-E time points. **c** Sample I-V curves of wild-type hTRPM4 and its alanine substitutions at the PI(4,5)P_2_-binding site recorded in whole-cell patches with 10 µM NC1 in the bath (extracellular). **d** Outward and inward current densities of NC1-activated wild-type hTRPM4 and its mutants recorded at ±100 mV in whole-cell patches with 10 µM NC1 in the bath (extracellular). Bars represent mean ± SEM of n=5 independent replicates (shown as dots). p-values were calculated using a two-sided Student’s t-test and are provided in the Source Data file. ** represents p < 0.01 and * represents p < 0.05.

To further support this conclusion, we mutated key positively charged PI(4,5)P_2_-interacting residues, R664A, K925A, and K928A, previously identified in the Ca²⁺/PI(4,5)P_2_-activated TRPM4 structure^36^. With surface expression levels comparable to the wild-type channel (Supplementary Fig. 4a), these mutations significantly impaired NC1 activation in whole-cell recordings, with much stronger effects on inward currents at negative potentials (Fig. 4c and 4d). Additionally, HeLa cells overexpressing these mutants were no longer sensitive to NC1-induced cell death, further validating the requirement of PI(4,5)P_2_ for NC1-mediated TRPM4 activation (Supplementary Fig. 4b).

The PI(4,5)P_2_ dependence of NC1-induced hTRPM4 activation was further confirmed by determining the structure of the channel bound to both PI(4,5)P_2_ and NC1 at 2.8 Å resolution (Fig. 5a and 5b, Supplementary Fig. 1b, Supplementary Table 1, and Methods). In this structure, hTRPM4 adopts an open conformation virtually identical to that of the Ca²⁺/PI(4,5)P_2_-activated channel (Fig. 5a). Compared to the structure with NC1 alone, the presence of PI(4,5)P_2_ enables the NC1-induced conformational changes within the S1-S4 domain to be effectively transmitted to the pore through the reformation of the critical F910/F935 contact, thereby leading to pore opening (Fig. 3c and 3d and Fig. 5c and 5d). This pore-opening movement is further coupled to large-scale rotational and upward swinging motions of the cytosolic domains via the TRP domain, as also observed in the Ca²⁺/PI(4,5)P_2_-activated structure^36^ (Fig. 5c and 5e). Interestingly, despite no exogenous Ca²⁺ being added, the canonical Ca²⁺-binding site within the S1-S4 domain is occupied by a Ca²⁺ ion in the NC1/PI(4,5)P_2_-bound structure (Fig. 5b). The stabilized, active conformation of S1-S4 induced by NC1 and PI(4,5)P_2_ likely increases the apparent Ca²⁺ affinity, allowing residual micromolar Ca²⁺ in nominally Ca²⁺-free conditions to bind. To confirm that Ca²⁺ is not required for channel opening in the NC1/PI(4,5)P_2_-bound structure, we also determined the structure of the complex prepared in the presence of EGTA (Supplementary Fig. 1c, Supplementary Table 1, and Methods). In this case, no bound Ca²⁺ is observed, yet the channel remains in the same open conformation, confirming that NC1 and PI(4,5)P_2_ are sufficient to activate the channel independently of Ca²⁺ (Supplementary Fig. 5a and 5b). Furthermore, the presence of Ca^2+^ does not appear to exert any potentiating effect on NC1 activation, as NC1/PI(4,5)P_2_-activated currents are comparable when recorded in both the absence and presence of Ca^2+^ (Supplementary Fig. 5c).

**Fig. 5:**
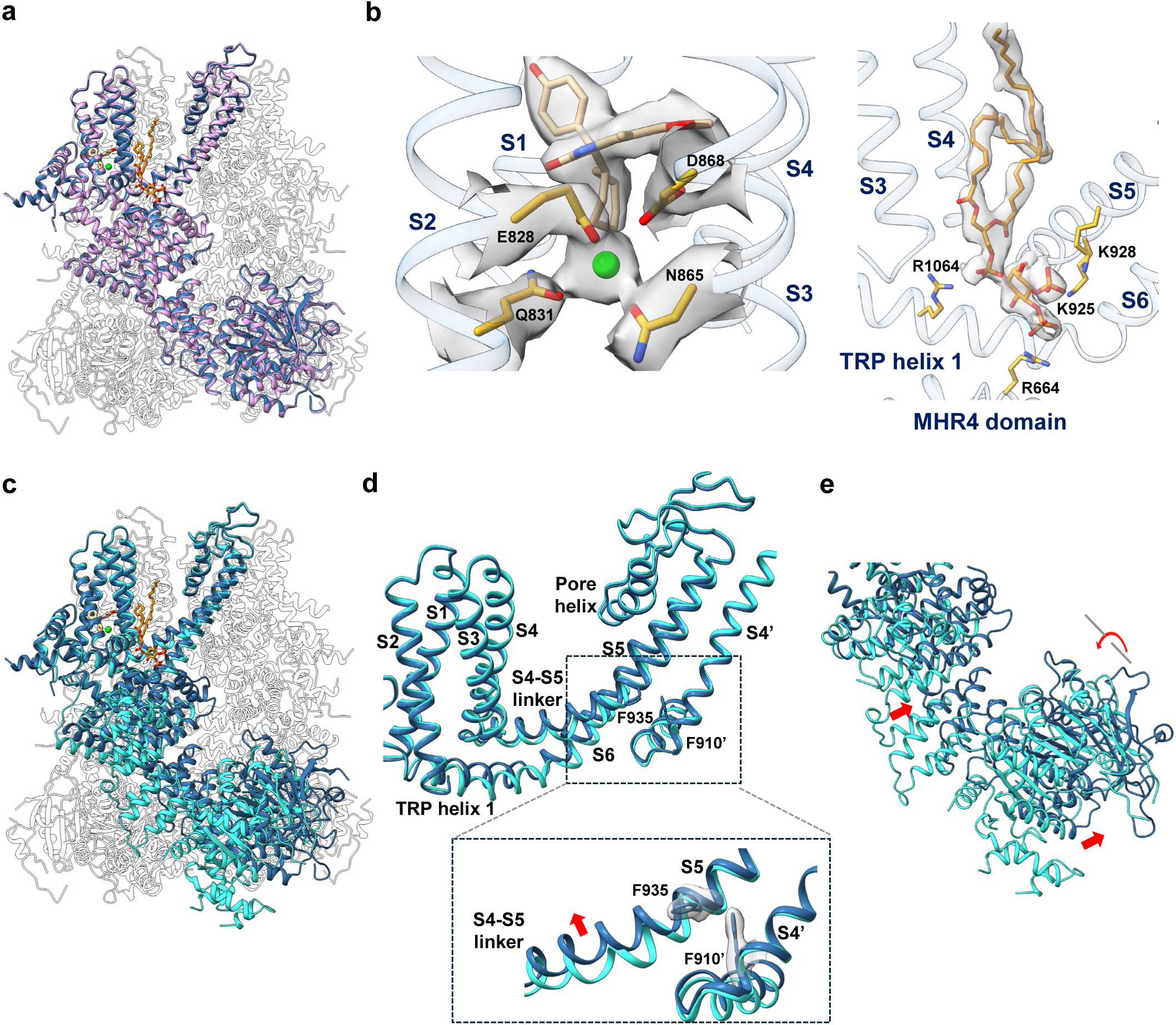
Structure of NC1/PI(4,5)P_2_-bound hTRPM4. **a** Superposition of the NC1/PI(4,5)P_2_-bound and Ca^2+^/PI(4,5)P_2_-bound (PDB:9MRT) hTRPM4 open structures with the front subunits highlighted in navy blue and pink, respectively. NC1 and PI(4,5)P_2_ are depicted in stick representation, and Ca^2+^ is shown as a green sphere. **b** Zoomed-in view of the ligand binding sites in the NC1/PI(4,5)P_2_-bound structure. Ca^2+^ is represented as a green sphere. NC1, PI(4,5)P_2,_ and ligand-interacting residues are shown in stick representation. Density (grey surface) is contoured at 0.48 for NC1, Ca^2+^ and Ca^2+^ interacting residues and at 0.45 for PI(4,5)P_2_ in ChimeraX. **c** Superposition of the NC1/PI(4,5)P_2_-bound open and NC1-bound hTRPM4 structures with the front subunits highlighted in navy blue and cyan, respectively. **d** Enlarged view of the conformational changes at the transmembrane region between the NC1/PI(4,5)P_2_-bound open (navy blue) and NC1-bound (cyan) structures. The S4 helix from the neighboring subunit (S4’) is also shown. The inset shows a close-up highlighting the coupling between F935 and neighboring F910’ in the NC1/PI(4,5)P_2_-bound structure that leads to the movement (red arrow) at the S4-S5 linker and S5 and channel opening. Density for F935 and F910’ is shown in a grey surface and contoured at 0.55 in ChimeraX. **e** Conformational changes at the cytosolic MHR domains between the NC1/PI(4,5)P_2_-bound (navy blue) and NC1-bound (cyan) structures. Red arrows mark the upward swing of the cytosolic MHR domains and the rotation of MHRs 1 and 2.

Together, these structural and functional results reveal several key insights into PI(4,5)P_2_-dependent NC1 activation: NC1 induces conformational changes in the S1-S4 domain similar to those triggered by Ca²⁺ without interfering with Ca²⁺ binding; Like Ca²⁺, NC1 alone stabilizes the channel in a desensitized state, requiring PI(4,5)P_2_ to promote channel opening; In the presence of PI(4,5)P_2_, both NC1 and Ca²⁺ can independently activate hTRPM4.

### Molecular determinants of NC1 specificity for human TRPM4

NC1 activates human but not mouse TRPM4^40^, despite the high sequence and structural homology between the two orthologs. This species-specific activation is particularly intriguing given the identical amino acid composition and conserved structure of the NC1-binding pocket. To identify the molecular determinants underlying this specificity, we used the cell death assay to screen various mutations in both human and mouse TRPM4 expressed in HeLa cells. Through a combination of chimeric constructs and point mutations (Supplementary Fig. 6, Supplementary Fig. 7, and associated legends), we identified three residues that potentially contribute to the selective activation of hTRPM4 by NC1: V904 on the S4 helix (corresponding to L900 in mTRPM4), P997 in the pore domain (R993 in mTRPM4), and R1064 on the TRP helix (S1060 in mTRPM4) (Fig. 6a).

**Fig. 6:**
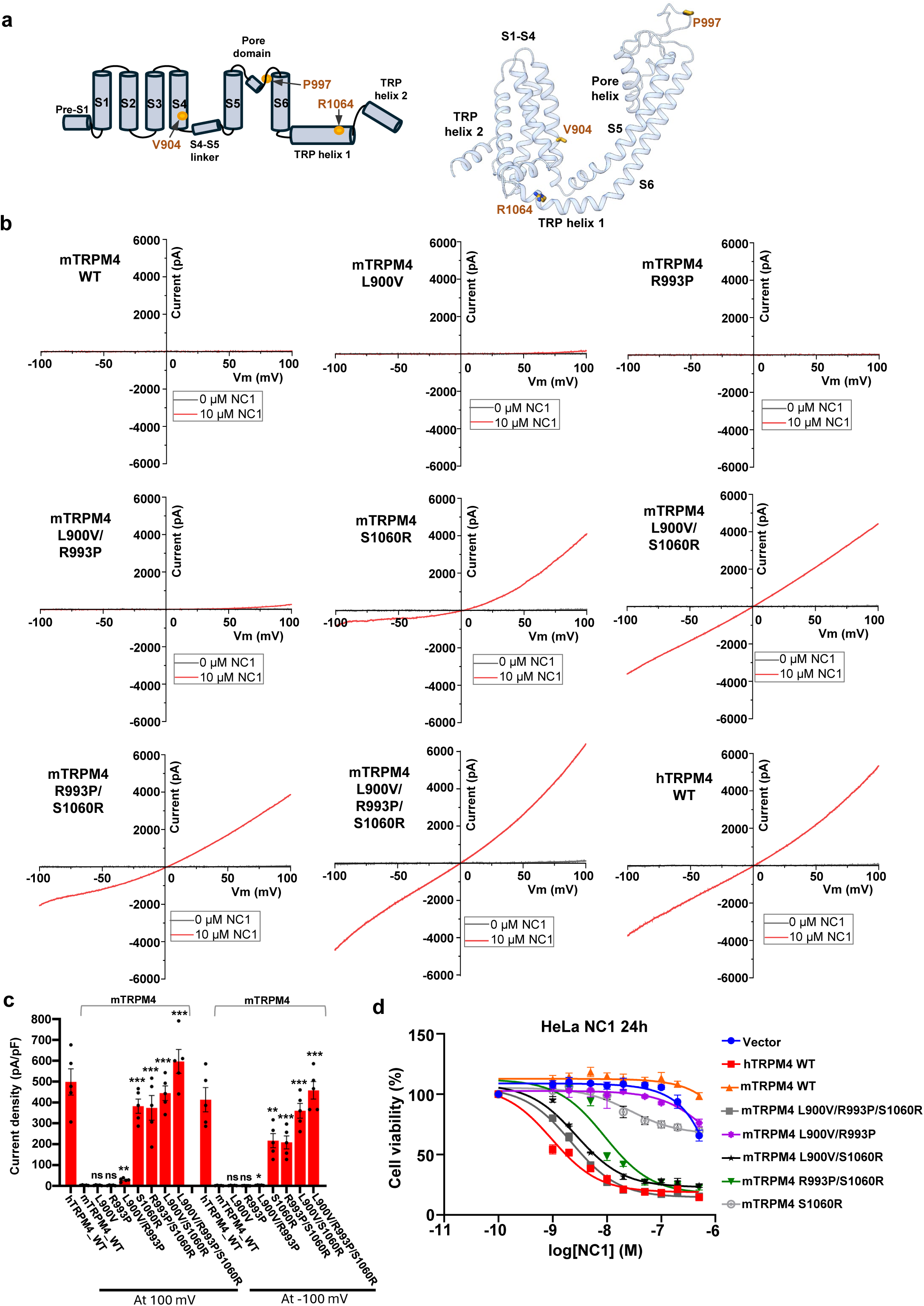
Molecular determinants of NC1 specificity for human TRPM4. **a** Topology (left) and structure (right) of the hTRPM4 TM domain with the three key residues important for NC1 specificity marked. **b** Sample I-V curves of wild-type mTRPM4 and its single, double, and triple mutants at the three key residues. Currents were recorded in whole-cell patches with 10 µM NC1 in the bath (extracellular). Current from NC1-activated wild-type hTRPM4 is shown in the last panel for comparison. **c** Outward and inward current densities of the wild-type mTRPM4 and its mutants recorded at ±100 mV in whole-cell patches with 10 µM NC1 in the bath (extracellular). Bars represent mean ± SEM of n=5 independent replicates (shown as dots). p-values were calculated using a two-sided Student’s t-test and are provided in the Source Data file. *** represents p < 0.001, ** represents p < 0.01 and * represents p < 0.05. Current densities of hTRPM4 are also graphed for comparison. **d** Dose-dependent NC1 cytotoxicity in stable HeLa cell lines expressing wild-type hTRPM4, wild-type mTRPM4, or mTRPM4 mutants. Data points represent mean ± SD of n = 3 independent replicates.

We next tested whether mouse TRPM4 could be converted into an NC1-sensitive channel by introducing single, double, and triple mutations in which the three residues (L900, R993, and S1060) were replaced with their human TRPM4 counterparts. Figure 6b and Figure 6c show the whole-cell patch-clamp recordings of NC1 activation in these mutants. Strikingly, the single S1060R mutation in mTRPM4 was sufficient to confer NC1 sensitivity; however, the resulting current was outwardly rectifying, with greater conductance at depolarized membrane potentials.

In contrast, mTRPM4 mutants bearing L900V and/or R993P substitutions remained insensitive to NC1 when tested individually or in combination. Notably, when these mutations were combined with S1060R, NC1 sensitivity was significantly enhanced. The mTRPM4 triple mutant (L900V/R993P/S1060R) displayed an NC1 activation profile nearly identical to that of hTRPM4. As these mutations do not target the Ca^2+^- or PI(4,5)P_2_-binding sites, they are not expected to affect Ca^2+^/PI(4,5)P_2_-mediated activation of mTRPM4. To test that, we measured Ca^2+^/PI(4,5)P_2_-activated currents for all of these mutants using inside-out patch recordings and compared their sensitivity to NC1/PI(4,5)P_2_ activation (Supplementary Fig. 8a and 8b). The results show that all mutants produced Ca^2+^/PI(4,5)P_2_-elicited currents comparable to those of wild-type mTRPM4, indicating similar channel expression levels between the mutants and the WT. However, only mutants containing the S1060R mutation could be partially (single and double mutants) or fully (triple mutant) activated by NC1/PI(4,5)P_2_, consistent with the observations from the whole-cell recordings. Complementing these gain-of-function experiments, we also tested loss-of-function mutations in hTRPM4 (Supplementary Fig. 8c and 8d). Consistent with the mTRPM4 mutagenesis findings, mutations at V904 and P997 did not affect NC1 activation of hTRPM4. In contrast, both the single R1064S mutation and the V904L/P997R/R1064S triple mutation significantly reduced NC1 activation, though not completely abolishing it.

In addition, we generated the HeLa stable cell lines expressing those mTRPM4 gain-of-function mutants and measured their viability in response to NC1 treatment (Fig. 6d). Consistent with our electrophysiology result, expressing mTRPM4 with L900V and R993P mutations has little effect on cell viability similar to control cells expressing WT mTRPM4 or empty vector; however, the expression of S1060R single mutant or S1060R-containing double or triple mutants progressively decreases the cell viability in response to NC1; cells expressing the L900V/R993P/S1060R triple mutant of mTRPM4 exhibit similar sensitivity to NC1-induced necrosis as hTRPM4-expressing cells.

### The structural basis of NC1 specificity

To gain structural insight into the NC1 specificity of TRPM4, we determined the cryo-EM structures of both the NC1-insensitive WT mTRPM4 and its NC1-activated L900V/R993P/S1060R triple mutant, each in the presence of NC1 (Supplementary Fig. 9, Supplementary Table 2, and Methods). As PI(4,5)P₂ is required for NC1 activation in hTRPM4, it was also included during mTRPM4 protein preparation (Methods).

The structure of WT mTRPM4 in the presence of NC1 and PI(4,5)P_2_ was resolved at 2.9 Å resolution (Fig. 7a). Surprisingly, NC1 still binds to mTRPM4 at the same pocket as in hTRPM4 with identical surrounding residues (Fig. 7b), despite failing to activate the channel. Relative to the apo mTRPM4 structure, NC1 binding induces similar local conformational changes within the S1-S4 domain, as observed in hTRPM4 (Supplementary Fig. 10a). However, unlike NC1/PI(4,5)P_2_-activated hTRPM4, where large swinging and rotational movements of the cytosolic domains are coupled to pore opening, the cytosolic domains of NC1/PI(4,5)P_2_-bound mTRPM4 closely resemble the apo closed conformation, with only a subtle rigid-body shift (Fig. 7a). Notably, three distinct conformational changes in the transmembrane region of NC1/PI(4,5)P_2_-bound mTRPM4 distinguish it from the NC1-sensitive hTRPM4 and likely underlie its lack of activation (Fig. 7c, 7d and 7e). Firstly, the entire S1-S4 domain moves laterally toward the center of the channel (Fig. 7c). Secondly, the central filter and surrounding pore helices exhibit no clear density, suggesting the filter becomes disordered upon NC1 binding (Fig. 7d) - a surprising finding, as this tightly packed central filter region remains steady during channel gating and is typically the best-resolved in TRPM4 structures. Thirdly, the S5 helix shifts inward toward the filter, while the pore-lining S6 helix moves outward, widening the intracellular gate (Fig. 7c and 7e).

**Fig. 7:**
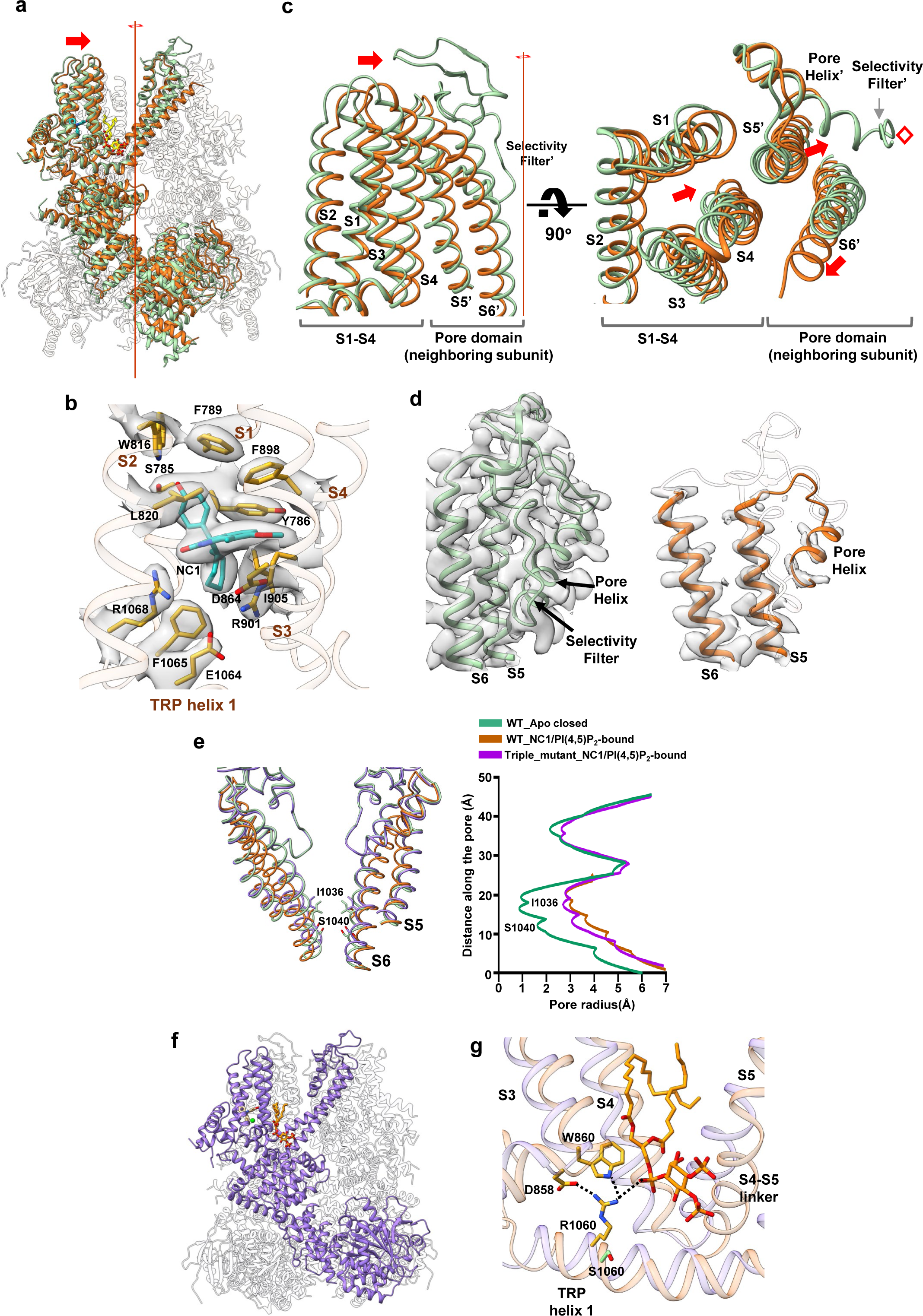
NC1 binding in mTRPM4. **a** Superposition of the NC1/PI(4,5)P_2_-bound (brown) and apo closed (PDB:6BCJ) (green) wild-type mTRPM4 structures along the pore axis, with the front subunits highlighted. NC1 (blue) and PI(4,5)P_2_ (yellow) are shown in stick representation. **b** Zoomed-in view of the NC1 binding site in mTRPM4. NC1 and its surrounding residues are shown in stick representation. Density for NC1 and surrounding residues (grey surface) is contoured at 0.23 in ChimeraX. **c** Conformational changes at S1-S4 and its neighboring pore domain upon NC1 binding to mTRPM4. The red line and red diamond mark the central 4-fold axis. Red arrows indicate the major movements in this region, namely the lateral movement of S1-S4 towards the central axis, the inward movement of the neighboring subunit S5 (S5’), and the outward movement of the neighboring subunit S6 (S6’). **d** Comparison of the density maps at the pore domain between the apo closed (PDB:6bcj) (green) and NC1/PI(4,5)P_2-_bound (brown) mTRPM4 structures. The disordered region in the pore domain of the NC1/PI(4,5)P_2-_bound structure is depicted as a semi-transparent cartoon. **e** Structural comparison of the ion conduction pore in the apo closed (green), NC1/PI(4,5)P_2_-bound inactivated (brown) wild-type mTRPM4 and in the NC1/PI(4,5)P_2_-bound open (purple) triple mutant (L900V/R993P/S1060R). Gating residues I1036 and S1040 are shown in stick representation. The front and back subunits were removed for clarity. The panel on the right depicts the pore radius along the central axis in these structures. The dotted line illustrates the disordered filter region in the NC1/PI(4,5)P_2_-bound wild-type mTRPM4. **f** Structure of the NC1/PI(4,5)P_2-_bound mTRPM4 triple mutant (L900V/R993P/S1060R) with the front subunit highlighted in purple. NC1 and PI(4,5)P_2_ are shown in stick representation and Ca^2+^ as a green sphere. **g** Structural comparison at S1060 region between NC1/PI(4,5)P_2-_bound wild-type mTRPM4 (light brown) and its triple mutant (L900V/R993P/S1060R, light purple). PI(4,5)P_2_ and key residues important for stabilizing S1-S4 in the mutant are shown in stick representation. The wild-type S1060 side chain is also shown for comparison.

The latter two structural changes appear to originate from the lateral movement of S1-S4. Due to tight S5-mediated packing between the S1-S4 domain and the neighboring subunit’s pore domain, this lateral shift of S1-S4 drives S5 inward, compressing the adjacent pore helix and filter toward the central axis (Fig. 7c). Unable to accommodate this compression, the filter likely undergoes a promiscuous movement and adopts a disordered, non-conductive conformation, leading to the collapse of the ion conduction pathway. The accompanying outward motion of S6 likely serves to relieve the spatial strain caused by the rearrangement of the central pore. Importantly, these conformational changes are solely induced by NC1 binding, as an NC1-bound mTRPM4 structure prepared without PI(4,5)P_2_ is identical to the NC1/PI(4,5)P_2_-bound form (Supplementary Fig. 9b, Supplementary Fig. 10b, Supplementary Table 2, and Methods). Thus, the insensitivity of mTRPM4 to NC1 activation is not due to a lack of drug binding, but rather to NC1-induced collapse of the selectivity filter, rendering the channel non-conductive, analogous to C-type inactivation observed in certain K⁺ channels^46^.

We next determined the structure of the NC1-sensitive mTRPM4 triple mutant (L900V/R993P/S1060R) in complex with NC1 and PI(4,5)P_2_ (Fig. 7f, Supplementary Fig. 9c, Supplementary Table 2, and Methods). Unlike WT mTRPM4, the mutant’s pore domain is well structured and adopts an open conformation (Fig. 7e), and its S1-S4 domain no longer undergoes a lateral movement. The overall structure of the mutant closely resembles that of the NC1/PI(4,5)P_2_-bound open hTRPM4 (Supplementary Fig. 10c), faithfully recapitulating the activation mechanism observed in the human channel. Given that the S1060R mutation in the TRP helix is the key determinant for conferring NC1 sensitivity, we focused our structural comparison on this region between NC1/PI(4,5)P_2_-bound WT and mutant mTRPM4. In the mutant, R1060 forms a network of inter-domain interactions absent in the WT channel (Fig. 7g). Specifically, R1060 mediates a cluster of salt bridges and hydrogen bonds involving D858 and W860 at the N-terminus of S3, as well as the glycerol phosphate head group of PI(4,5)P_2_. These interactions serve to stabilize the S1-S4 domain, preventing the inward lateral movement observed in NC1-bound WT mTRPM4 that disrupts the pore. Notably, this arginine residue, crucial for NC1 sensitivity, is conserved in human and primate TRPM4 orthologs but is replaced by a serine in most other mammalian species.

## Discussion

The small molecule NC1 activates hTRPM4, triggering Na⁺ influx and leading to necrotic cell death via sodium overload. Through structural and functional analyses, we elucidated the molecular mechanism underlying NC1-mediated activation of hTRPM4. NC1 binds near the Ca²⁺ activation site within the S1-S4 domain and induces local conformational changes that mirror those caused by Ca²⁺. Like Ca²⁺, NC1 requires membrane PI(4,5)P₂ to stabilize the open channel conformation. Thus, NC1 acts as a non-competitive surrogate of the native ligand. NC1 engages in extensive hydrophobic and polar interactions with surrounding residues, resulting in high-affinity binding. Its binding site, deeply buried within the protein, likely contributes to its slow activation kinetics and poor washout. The NC1 binding site is similar to that of the TRPM4 modulator deacetyl bisacodyl (DAB)^47^ and to some modulator-binding pockets in TRPM3 (e.g., primidone and CIM 0216)^48,49^ and TRPM8 (e.g., N-(3-aminopropyl)-2-[(3-methylphenyl)methoxy]-N-[(thiophen-2-yl)methyl]benzamide (AMTB), cryosim-3 (C3), icilin, and the menthol analog WS-12)^50–52^.

NC1 selectively activates human TRPM4 but not most other mammalian TRPM4 orthologs. Through comparative studies of human and mouse TRPM4, we determined that the species specificity of NC1 activation is not due to a failure in drug binding. Instead, in mTRPM4, NC1 binding destabilizes the selectivity filter, rendering the channel non-conductive, akin to C-type inactivation observed in certain K⁺ channels. We identified R1064, located on the TRP helix of hTRPM4, as a key determinant of NC1 specificity. This arginine mediates a network of interdomain interactions that stabilize the S1-S4 domain’s lateral position within the membrane. In TRPM4s from mice and most other mammals, a serine occupies this position. The absence of arginine-mediated stabilization permits a lateral shift of the S1-S4 domain upon NC1 binding, which collapses the selectivity filter and inactivates the channel. Interestingly, akin to NC1, the TRPM4 modulator CBA was shown to inhibit the human but not the mouse channel^34^ and does not exert any effect on other TRPM channel subfamily members^32^. However, the mechanistic details that confer TRPM4 species-specificity to this compound have not been thoroughly explored.

The mechanistic insights into NC1 activation of hTRPM4 and its remarkable specificity revealed in this study establish a structural framework for understanding how small molecules can selectively modulate TRPM4 function. The defined NC1-binding mode provides an essential molecular basis for rational drug design aimed at exploiting TRPM4-mediated necrosis as a potential therapeutic avenue for cancer treatment.

The discovery that NC1 binds to an identical site in both human and mouse TRPM4 orthologs but induces markedly different functional outcomes emphasizes the structural subtleties governing allosteric coupling and species-dependent channel gating. Given the widespread reliance on murine and other mammalian models in preclinical studies, our findings underscore the necessity of validating compound efficacy and mechanism in human-derived systems to ensure accurate interpretation of pharmacological responses. This cross-species discrepancy also serves as a cautionary example for drug development pipelines targeting ion channels, in which even small structural variations can yield divergent functional consequences. Collectively, our results provide a comprehensive molecular foundation for future high-throughput screening, structure-guided optimization, and in vivo evaluation of TRPM4 modulators.

## Methods

### Protein expression and purification

Full-length wild-type human TRPM4 (UniProtKB: Q8TD43) containing an N-terminal FLAG-tag (DYKDDDDK) and full-length wild-type or L900V;R993P;S1060R mouse TRPM4 (UniprotKB: Q7TN37) containing a C-terminal FLAG-tag were cloned into a pEZT-BM plasmid^53^. *E.coli* DH10Bac cells were used to synthesize the bacmid that was employed in baculovirus production in Sf9 cells (Thermo Fisher Scientific) using Cellfectin II reagent (Thermo Fisher Scientific). For protein expression, HEK293S GnTI^-^ cells (American Type Culture Collection (ATCC), CRL-3022) grown in suspension to a density of 3×10^6^ cells/mL were infected with P3 viruses at a ratio of 1:40 (virus:cell, v/v) and supplemented with 10 mM sodium butyrate to boost protein expression. The cells were then incubated at 37 °C for 48 h before being harvested by centrifugation (5000 g, 15 min, 4 °C). The cell pellet was resuspended in lysis buffer (50 mM Tris-HCl pH 7.5, 150 mM NaCl) supplemented with protease inhibitors (0.5 μg/ml pepstatin, 2 μg/ml leupeptin, 1 μg/ml aprotinin, and 1 mM PMSF) and homogenized by sonication. TRPM4 was then extracted with 1 % (w/v) lauryl maltose neopentyl glycol (LMNG, Anatrace) in gentle agitation for 2 h at 4 °C. The supernatant was subsequently collected by centrifugation (40000 g, 40 min, 4 °C) and incubated with Anti-DYKDDDDK G1 Affinity Resin (GenScript) for 1 h and 30 min at 4 °C in gentle agitation. The resin was then washed with Buffer A (50 mM Tris-HCl pH 7.5, 150 mM NaCl, 0.01 % LMNG) and elution was performed by incubating for 45 min at room temperature in gentle agitation with Buffer B (50 mM Tris-HCl pH 7.5, 150 mM NaCl, 0.01 % LMNG, 0.2 mg/mL FLAG peptide). The eluate was then concentrated and further purified by size-exclusion chromatography in Buffer C (50 mM Tris-HCl pH 7.5, 150 mM NaCl, 0.0035 % LMNG) on a Superose 6 Increase 10/300 GL (GE Healthcare). For the Ca^2+^ free samples, 2 mM EGTA was also included in Buffer C and for the samples requiring PI(4,5)P_2_, 50 µM Brain PI(4,5)P_2_ (L-α-phosphatidylinositol-4,5-bisphosphate (Brain, Porcine) (ammonium salt), Avanti) was added to the buffer B during the elution step and further used to supplement the concentrated eluate before size-exclusion chromatography.

### Cryo-EM sample preparation and data acquisition

hTRPM4 and mTRPM4 samples were concentrated to ∼7-8 mg/mL and supplemented with 0.3 mM Necrocide 1 ((S’)-3-cycloheptyl-3-(4-hydroxyphenyl)-6-methoxy-7-methyl-1,3-dihydroindole-2-one) (NC1). 1 mM EGTA was also added when required. Supplemented samples were incubated for 1 h on ice before vitrification. 4 µL was then applied to a glow-discharged Quantifoil R1.2/1.3 300-mesh gold holey carbon grid (Quantifoil, Micro Tools GmbH, Germany), blotted for 3.5 s under 100 % humidity at 12 °C and plunged into liquid ethane using a Mark IV Vitrobot (FEI).

All data collections were performed using SerialEM with a nominal defocus range set from -0.9 to -2.2 μm and each movie was recorded with a total dose of 60 e^-^/Å^2^.

For the dataset of hTRPM4 containing NC1 and EGTA, movies were acquired on a Titan Krios microscope (FEI) operated at 300 kV with a Falcon4i direct electron detector camera (Thermo Fisher Scientific) at 0.738 Å pixel size.

For the dataset of hTRPM4 containing NC1 and PI(4,5)P_2_, movies were acquired on a Titan Krios microscope (FEI) operated at 300 kV with a K3 direct electron detector camera (Gatan) using the CDS (Correlated Double Sampling) mode with a super-resolution pixel size of 0.4172 Å.

For the datasets of hTRPM4 containing NC1, PI(4,5)P_2_ and EGTA and mTRPM4 wild-type containing NC1 and PI(4,5)P_2_, movies were acquired on a Titan Krios microscope (FEI) operated at 300 kV with a K3 direct electron detector camera (Gatan) using the CDS (Correlated Double Sampling) mode with a super-resolution pixel size of 0.4285 Å.

For the datasets of mTRPM4 wild-type containing NC1 and mTRPM4 L900V/R993P/S1060R triple mutant containing NC1 and PI(4,5)P_2_, movies were acquired on a Titan Krios microscope (FEI) operated at 300 kV with a Falcon4i direct electron detector camera (Thermo Fisher Scientific) at 0.735 Å pixel size.

### Cryo-EM data processing

The workflow for data processing is summarized in Supplementary Figure 1 and Supplementary Figure 9. Data processing was performed using cryoSPARC (versions 4.3.1 or 4.4.1)^54^ following the general scheme described below with some modifications between the datasets. For the dataset of hTRPM4 containing NC1 and PI(4,5)P_2_, RELION (version 3.1)^55,56^ was also employed in the processing workflow.

For the dataset of hTRPM4 containing NC1 and PI(4,5)P_2_, movies were motion-corrected and dose-weighted using MotionCor2^57^ and CTF estimation was performed using GCTF^58^. Micrographs were manually inspected to remove those with bad defocus values and ice contamination. Particles were then picked using Laplacian-of-Gaussian (LoG) and extracted in RELION. Extracted particles were subjected to multiple rounds of 2D classification and classes displaying clear features of the TRPM4 channel were used for subsequent 3D classification. The particles from the best-resolving 3D classes along with the respective 3D volumes were then imported into cryoSPARC and used for 3D heterogeneous refinement and 3D classification without alignment to remove junk particles and differentiate channel conformations. The best-resolving 3D classes were refined using Non-Uniform refinement with imposed C4 symmetry^59^.

For all other datasets, movies were subjected to patch motion correction and subsequent patch CTF estimation in cryoSPARC. The resulting micrographs were manually curated and those exhibiting drift, poor CTF estimation and outliers in defocus values and ice thickness were excluded from further processing (individually assessed for each parameter relative to the overall distribution). An initial round of particle picking was carried out with blob picker. Particles were then extracted and subjected to one round of 2D classification. Classes displaying clear features of the TRPM4 channel were selected and used either directly for subsequent processing or to re-pick particles with template picker. Additional rounds of 2D classification were further performed and particles from selected classes were used to obtain an initial 3D reconstruction with Ab-Initio. 3D heterogeneous refinement was then employed to remove junk particles, and the resulting particles were subjected to 3D classification without alignment to differentiate channel conformations. The best-resolving 3D classes were refined using Non-Uniform refinement with imposed C4 symmetry^59^.

Map resolutions were reported according to the gold-standard Fourier shell correlation (FSC) using the 0.143 criterion^60^. Local resolutions and angular distributions were estimated in cryoSPARC.

### Model building

Initial models were obtained using a combination of ModelAngelo^61^ and previously available hTRPM4 (PDBs: 9MTA and 9MRT) and mTRPM4 (PDB:6BCJ) structures. The models were then manually adjusted in Coot^62^ and refined against the respective maps in PHENIX^63^. Ligand restraints CIF files were generated using the Grade2 Web Server^64^. The geometry statistics of the models were obtained using MolProbity^65^. All the structural figures were prepared using UCSF ChimeraX^66,67^.

### Electrophysiology

Wild-type human and mouse TRPM4, along with the respective mutants, were cloned into a pEGFP-N1 plasmid (Clontech). Mutants were generated by site-directed mutagenesis using the QuickChange method and verified by sequencing. 1 µg of plasmid was transfected into HEK293 cells using Lipofectamine 2000 (Life Technology). 48 h after transfection, cells were dissociated by trypsin treatment and kept in a complete serum-containing medium before being re-plated onto 35-mm tissue culture dishes and incubated in a tissue culture incubator until recording.

Channel currents were recorded in whole cell or excised inside-out configuration. The long-chain native Brain PI(4,5)P_2_ employed in the structural studies is insoluble in water and forms liposomes in the recording solutions, making it difficult to fuse into the patch membrane. Therefore, the water-soluble short-chain synthetic PI(4,5)P_2_ diC8 (Phosphatidylinositol 4,5-bisphosphate diC8, Echelon Bioscience) was used in the functional experiments as needed. For inside-out patches, the standard bath solution (cytosolic side) contained (in mM): 145 caesium methanesulfonate (Cs–MS), 5 NaCl, 1 MgCl_2_, 0.3 CaCl_2_, 10 HEPES buffered with Tris, pH 7.4. 10 µM NC1 and/or 10 µM PI(4,5)P_2_ diC8 were added to the bath solution when required. For the calcium-free condition, 0.5 mM EGTA was added to the bath solution without CaCl_2_. The pipette solution (extracellular side) contained (in mM): 140 Na–MS, 1 MgCl_2_, 5 CaCl_2_, 10 HEPES buffered with Tris, pH 7.4. For whole cell current recordings, the pipette solution contained (in mM): Cs-MS 140, CsCl 4, MgCl_2_ 1, EGTA 1 or 10, HEPES 10, buffered with Tris, pH 7.4. Bath solution contained (in mM): Na-MS 140, NaCl 5, MgCl_2_ 1, CaCl_2_ 1, HEPES 10, buffered with Tris, pH=7.4. 10 µM NC1 was added to the bath solution as needed. Patch pipettes were pulled from borosilicate glass (Harvard Apparatus) and heat polished to a resistance of 3–5 MΩ. After the patch pipette was attached to the cell membrane, a giga seal (> 10 GΩ) was formed by gentle suction. The inside-out configuration was formed by pulling the pipette away from the cell and the pipette tip was exposed to air for a short time in some cases. The holding potential was set to 0 mV. The current and voltage relationship (I–V curve) was obtained using voltage pulses ramped from – 100 to + 100 mV over an 800 ms duration every 2 seconds. Data was acquired using an AxoPatch 200B amplifier (Molecular Devices) and a low-pass analog filter set to 1 kHz. The current signal was sampled at a rate of 20 kHz using a Digidata 1322A digitizer (Molecular Devices) and further analyzed with pClamp 11 software (Molecular Devices). Sample traces for the I–V curves of macroscopic currents shown were obtained from recordings on the same patch. Bars represent mean ± SEM of five measurements from different patches (n = 5 independent biological replicates).

### Cell death assay

Cell death was assessed in HeLa cells transiently expressing EGFP-tagged human or mouse TRPM4 by monitoring the morphological change of transfected cells using microscopy, as described previously^40^. Briefly, HeLa cells were seeded onto glass coverslips and grown to 30–40% confluency before transfection. 24 hours after TRPM4-EGFP transfection, cells were subjected to 1 µM NC1 treatment. Fluorescence images were taken before and after the indicated period of treatment. The morphological changes of cells with green fluorescence were monitored using microscopy, and over 50 cells expressing TRPM4 were analyzed in each group.

### Cell viability assay

Cell viability was assessed in HeLa cells stably expressing human or mouse TRPM4, using the CellTiter-Glo (CTG) assay as described previously^40^. Briefly, cells were seeded in a 96-well plate at a density of 1×10^4^ cells per well and allowed to adhere overnight. After treatment with various experimental conditions, the intracellular ATP was measured using the CTG luminescent assay kit (Promega, G7570) according to the manufacturer’s instructions. Briefly, 100 μL of CTG reagent was added to each well, and the plate was gently shaken for 10 minutes to induce cell lysis and release of intracellular ATP. The luminescence signal, proportional to the amount of ATP present and indicative of cell viability, was measured using a multi-well plate reader. All experiments were repeated three times, and the results were presented as mean ± SD.

### Cell surface biotinylation assay and western blot

Surface protein biotinylation was employed to assess the expression levels of wild-type hTRPM4 and respective PI(4,5)P_2_ binding site mutants used in the electrophysiology experiments. The assay was performed using the Cell Surface Biotinylation and Isolation Kit (Pierce, A44390) according to the manufacturer’s instructions. Briefly, transfected HEK293 cells were washed with PBS and incubated with Sulfo-NHS-SS-Biotin labeling solution for 30 min at 4 °C. The cells were then washed three times with ice-cold TBS to quench and remove the non-reacted biotinylation reagent, scraped, and collected by centrifugation (500 g, 3 min). Cell lysis was carried out using the supplied lysis buffer supplemented with protease inhibitors, and clarified lysates were collected by centrifugation (15000 g, 5 min, 4°C) and incubated with NeutrAvidin Agarose beads for 30 min at room temperature using gentle rotation. After washing, the bound biotinylated surface proteins were eluted by incubating the beads with the supplied elution buffer supplemented with 10 mM DTT for 30 min at room temperature using gentle rotation. Clarified lysates and eluted proteins were separated by SDS-PAGE and transferred to PVDF membranes for immunoblotting. hTRPM4 was probed using the anti-GFP-tag rabbit monoclonal antibody (Thermo Fisher Scientific, G10362). The surface protein CD44 was used as a loading control and probed with the Anti-CD44 rabbit polyclonal antibody (Abcam, ab157107).

## Data availability

The cryo-EM density maps of the human TRPM4 have been deposited in the Electron Microscopy Data Bank (EMDB) under accession numbers EMD-73526 (NC1-bound in the presence of EGTA), EMD-73527 (NC1/PI(4,5)P_2_-bound), and EMD-73528 (NC1/PI(4,5)P_2_-bound in the presence of EGTA). Atomic coordinates have been deposited in the Protein Data Bank (PDB) under accession numbers 9YVK (NC1-bound in the presence of EGTA), 9YVL (NC1/PI(4,5)P_2_-bound), and 9YVM (NC1/PI(4,5)P_2_-bound in the presence of EGTA).

The cryo-EM density maps of the mouse TRPM4 have been deposited in the Electron Microscopy Data Bank (EMDB) under accession numbers EMD-73530 (NC1/PI(4,5)P_2_-bound wild-type mTRPM4), EMD-73529 (NC1-bound wild-type mTRPM4), and EMD-73531 (NC1/PI(4,5)P_2_-bound L900V/R993P/S1060R triple mutant mTRPM4). Atomic coordinates have been deposited in the Protein Data Bank (PDB) under accession numbers 9YVO (NC1/PI(4,5)P_2_-bound wild-type mTRPM4), 9YVN (NC1-bound wild-type mTRPM4), and 9YVP (NC1/PI(4,5)P_2_-bound L900V/R993P/S1060R triple mutant mTRPM4).

All other data and materials supporting the findings of this study can be obtained from the corresponding authors upon reasonable request.

## Acknowledgements

Single particle cryo-EM data were collected at the University of Texas Southwestern Medical Center Cryo-EM Facility (CEMF) funded by the CPRIT Core Facility Support Award RP170644 and at Howard Hughes Medical Institute Janelia Cryo-EM Facility. Cryo-EM sample grids were prepared at the Structural Biology Laboratory at UT Southwestern Medical Center partially supported by grant RP170644 from CPRIT. This work was supported in part by the Howard Hughes Medical Institute (to Y.J.) and by grant from the National Institute of Health (R35GM140892 to Y.J.), and by grants from MOST (Ministry of Science and Technology of the People’s Republic of China) (2023YFA0914900 to Q.Z.), NSFC (National Natural Science Foundation of China) (W2511018, 32530055 to Q.Z., 82504119 to W.F.). This article is subject to HHMI’s Open Access to Publications policy. HHMI lab heads have previously granted a nonexclusive CC BY 4.0 license to the public and a sublicensable license to HHMI in their research articles. Pursuant to those licenses, the author-accepted manuscript of this article can be made freely available under a CC BY 4.0 license immediately upon publication.

## Author Contributions

C.T-D. prepared the purified protein samples, performed data acquisition, image processing, and structure determination; W.F. established the cell death assay system, led the functional characterization, and identified the key residues underlying the species-specific response of TRPM4 to NC1; W.Z. performed electrophysiology recording; J.W., X.J., and Z.Z assisted with mutagenesis and cell death assay; Y.J. and Q.Z. supervised the work. All authors participated in research design, data analysis, discussion, and manuscript preparation.

