## Supplementary figures and tables for "Structural mechanism of Necrocide 1 activation of human TRPM4 that triggers necrosis by sodium overload"

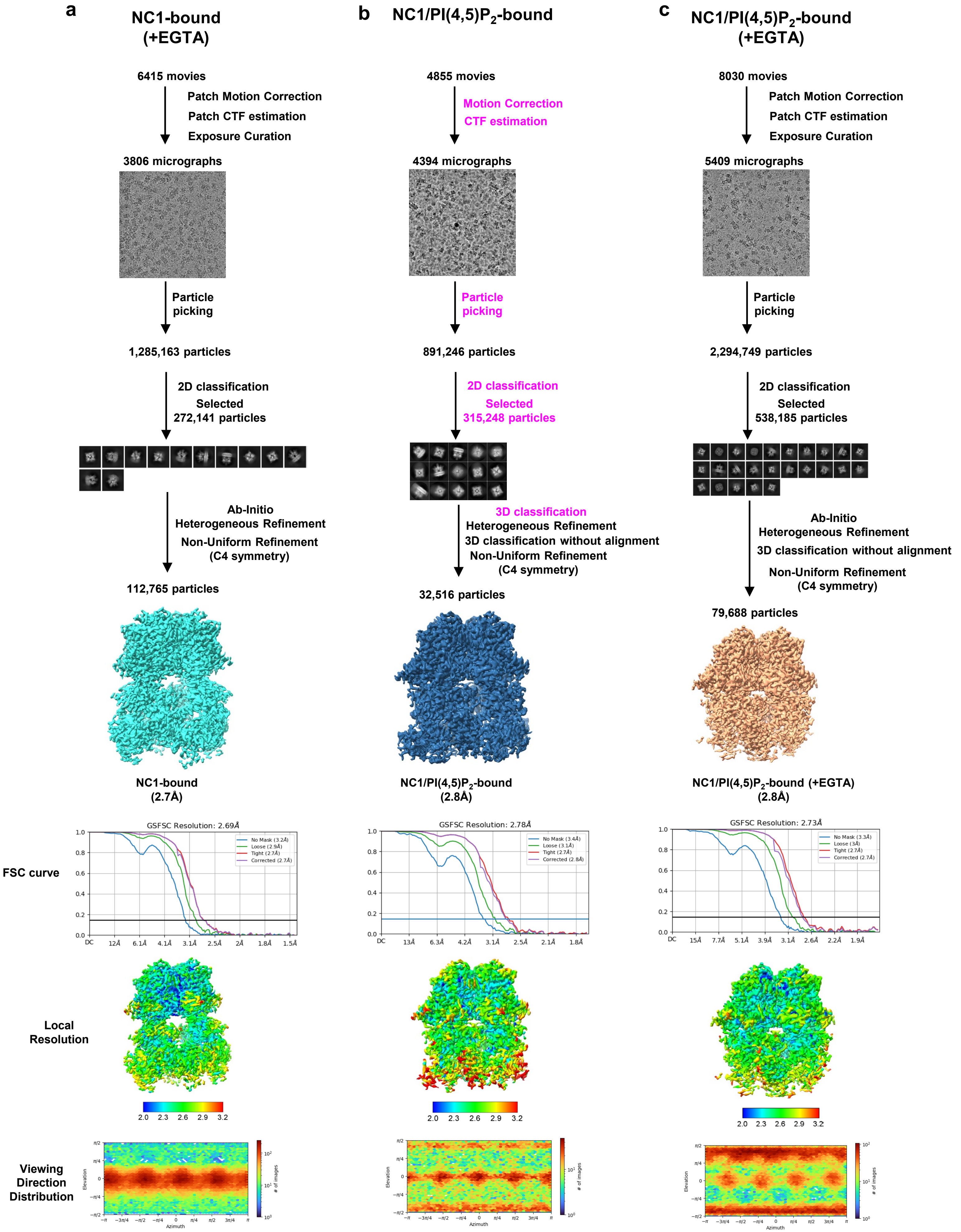


**Supplementary Fig. 1: Cryo-EM data processing scheme for the NC1-bound hTRPM4 samples.** Steps colored in purple were performed in RELION instead of CryoSPARC.

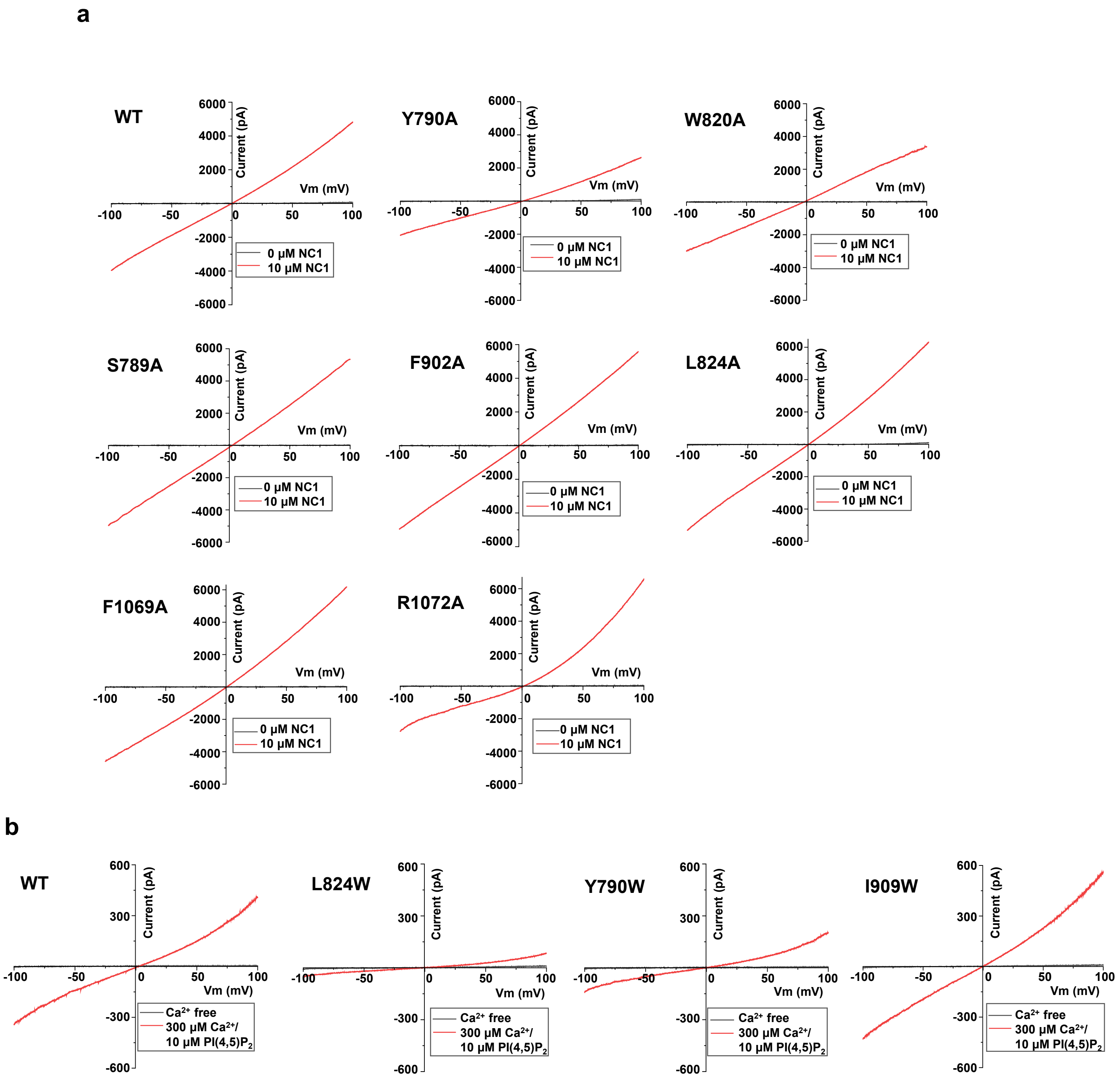


**Supplementary Fig. 2: Electrophysiology of hTRPM4 and its mutants at the NC1-binding site.** **a** Sample I-V curves of the wild-type hTRPM4 and its alanine substitutions at the NC1-binding site recorded in the whole-cell patches with 10 μM NC1 in the bath (extracellular). **b** Sample I-V curves of the wild-type hTRPM4 and its tryptophan substitutions at the NC1-binding site recorded in the inside-out patches with 300 μM $Ca^{2+}$ and 10 μM $PI(4,5)P_2$ diC8 in the bath (cytosolic).

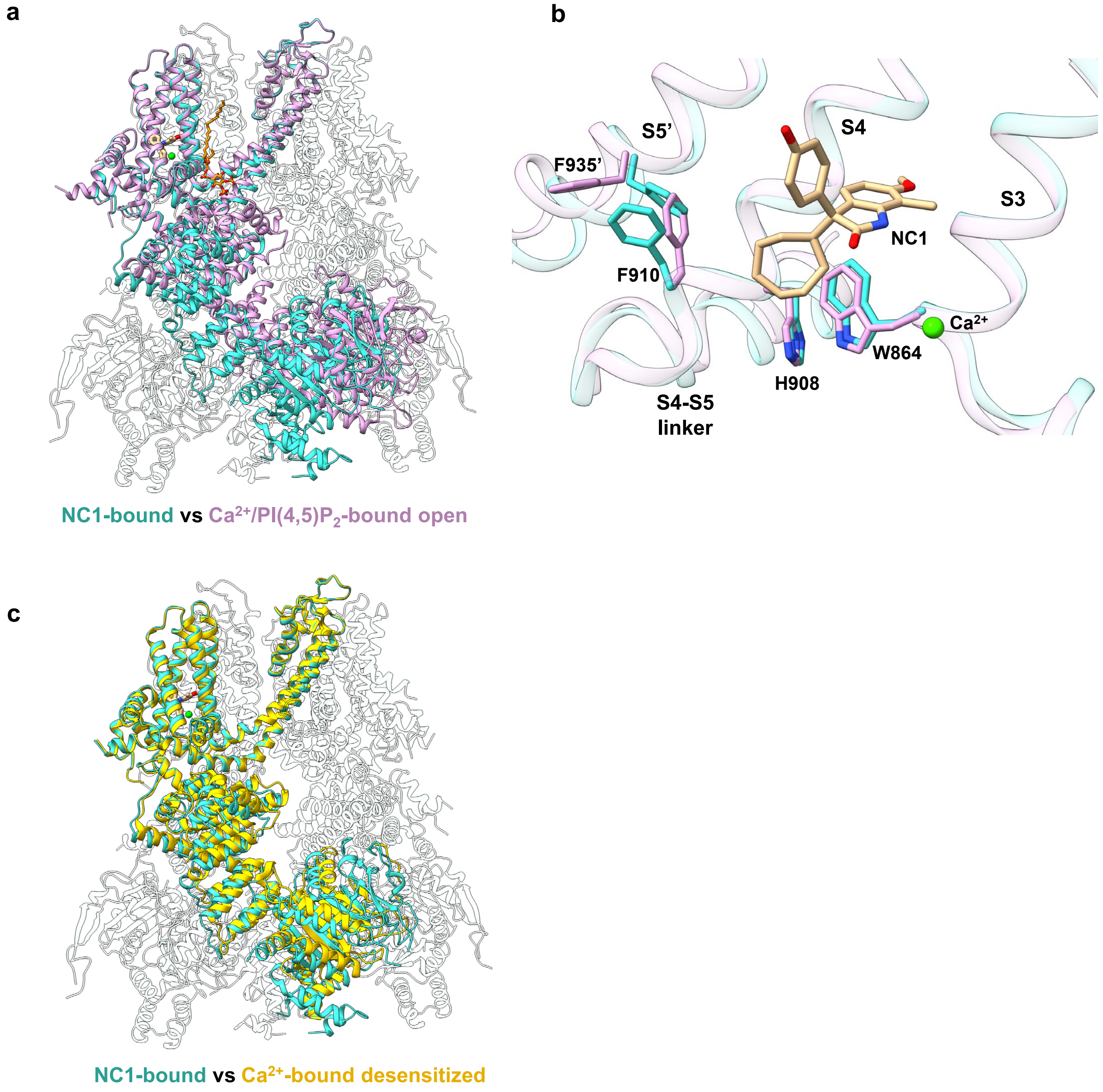


**Supplementary Fig. 3: Structural comparison of hTRPM4 in NC1-bound, $Ca^{2+}$-bound desensitized, and $Ca^{2+}$/PI(4,5)$P_2$-bound open states. a** Superposition of the hTRPM4 structures in NC1-bound (cyan) and $Ca^{2+}$/PI(4,5)$P_2$-bound open (pink) states with the front subunits highlighted. **b** Zoomed-in view of the superposition shown in **a** at the S1-S4 domain (only S3 and S4 helices are displayed) and the neighboring S5 (labeled with single quotation marks). Key residues for local conformational changes within S1-S4 upon TRPM4 activation are shown in stick representation. NC1-bound S1-S4 (light cyan) adopts the same structure as that in $Ca^{2+}$/PI(4,5)$P_2$-activated TRPM4 (light pink) but fails to induce any conformational change at the neighboring S5 for channel opening. **c** Superposition of the hTRPM4 structures in NC1-bound (cyan) and $Ca^{2+}$-bound desensitized (yellow) states with the front subunits highlighted. Both structures are highly similar and represent a desensitized conformation.

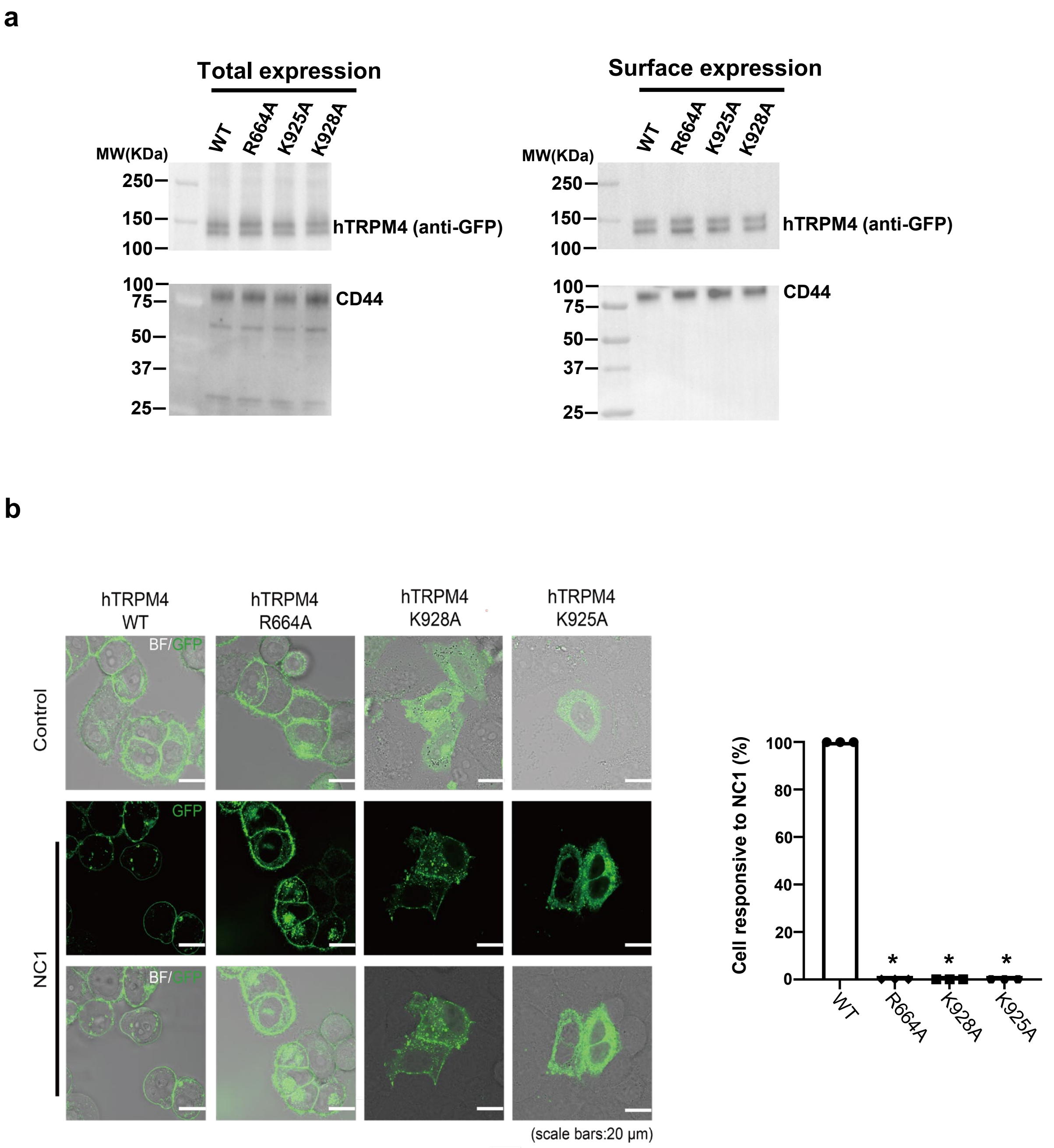


**Supplementary Fig. 4: Effect of hTRPM4 PI(4,5)$P_2$ binding site mutants on NC1-induced channel activation. a** Expression levels of hTRPM4 WT and PI(4,5)$P_2$-site mutants in HEK293 cells. Cell surface proteins were biotinylated and pulled down using NeutrAvidin Agarose beads. The cell surface glycoprotein CD44 was used as a loading control. **b** Effect of PI(4,5)$P_2$-binding site mutations on NC1-induced cell death in hTRPM4-expressing HeLa cells. Shown are sample images of WT or mutant hTRPM4-expressing HeLa cells 24 hours after NC1 treatment. The responsiveness of HeLa cells to NC1-induced cell death is quantified in the bar graph on the right. 20 cells were imaged in each replicate before and after NC1 treatment. Cells that present morphology changes after the treatment (“ballon-like” cell swelling) were counted as being responsive to NC1. All 20 hTRPM4 WT-expressing cells were responsive in all replicates and all mutant-expressing cells were non-responsive. Bars represent mean ± SEM of n = 3 independent replicates (shown as dots). p-values were calculated using a Kruskal-Wallis test and are provided in the Source Data file. * represents $p < 0.05$.

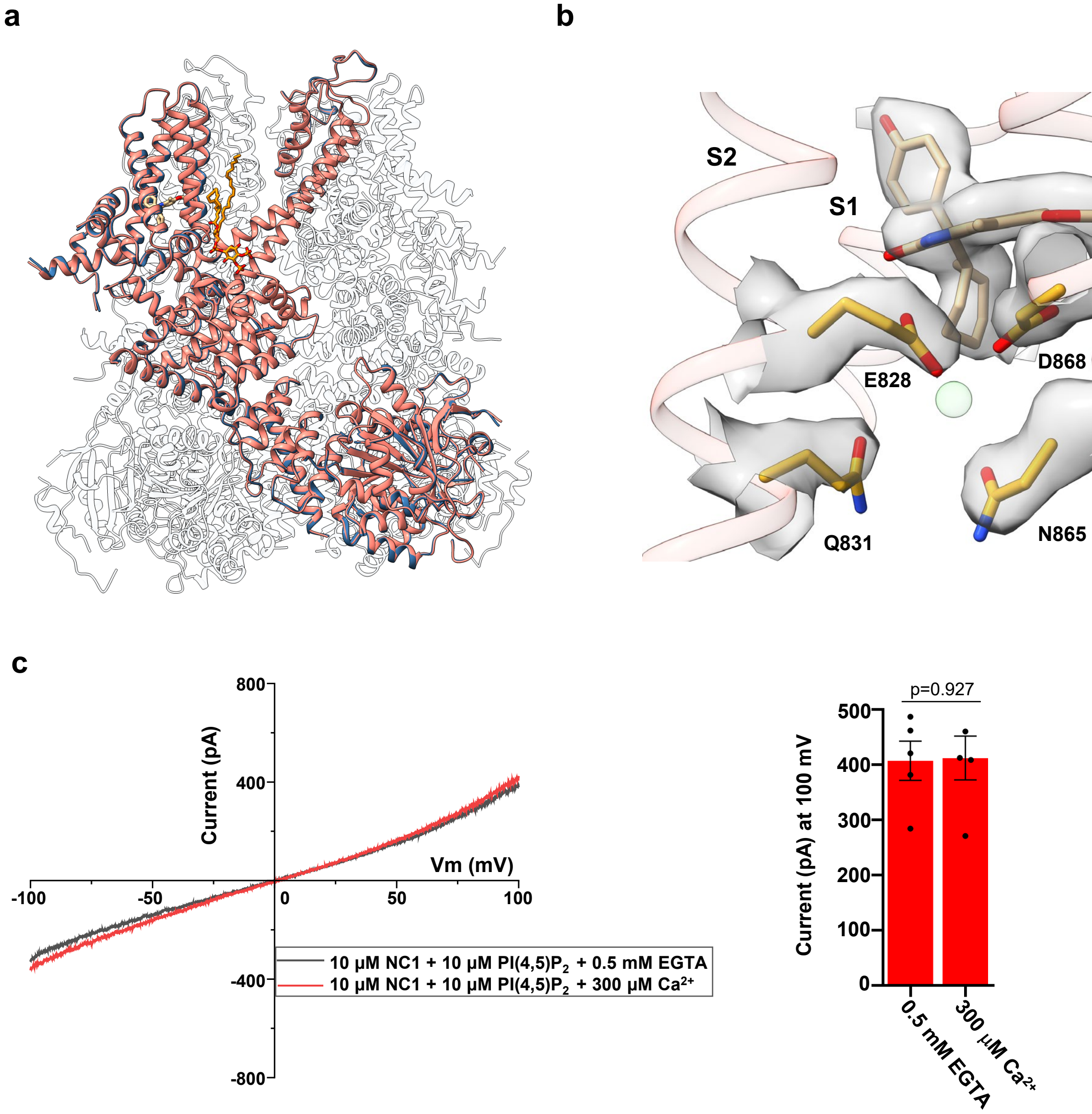


**Supplementary Fig. 5: Comparison of NC1/PI(4,5)$P_2$-bound hTRPM4 structures determined in the presence or absence of EGTA. a** Superposition of the NC1/PI(4,5)$P_2$-bound hTRPM4 structures determined in the presence (salmon) or absence (navy blue) of EGTA, with the front subunits highlighted. The two structures are virtually identical. **b** Zoomed-in view of the ligand-binding pocket within the S1-S4 domain of the NC1/PI(4,5)$P_2$-bound hTRPM4 structure obtained in the presence of EGTA. NC1 and key residues involved in $Ca^{2+}$ coordination are shown in stick representation, with the respective densities (grey surface) contoured at 0.35 in ChimeraX. No $Ca^{2+}$ density was observed at its activation site (illustrated with a light green sphere), and the channel remains in the open state. **c** Sample I-V curves from an inside-out patch of NC1/PI(4,5)$P_2$-activated hTRPM4 with 10 μM PI(4,5)$P_2$ diC8 and 10 μM NC1 in the bath (cytosolic) in the absence (with EGTA) or presence of $Ca^{2+}$. The bar graph on the right displays the outward currents recorded at +100 mV in the inside-out patches with 10 μM NC1, 10 μM PI(4,5)$P_2$ di-C8, and either 0.5 mM EGTA or 300 μM $Ca^{2+}$ in the bath (cytosolic). Bars represent mean ± SEM of n=5 independent replicates (shown as dots). p-value was calculated using a two-sided Student's t-test.

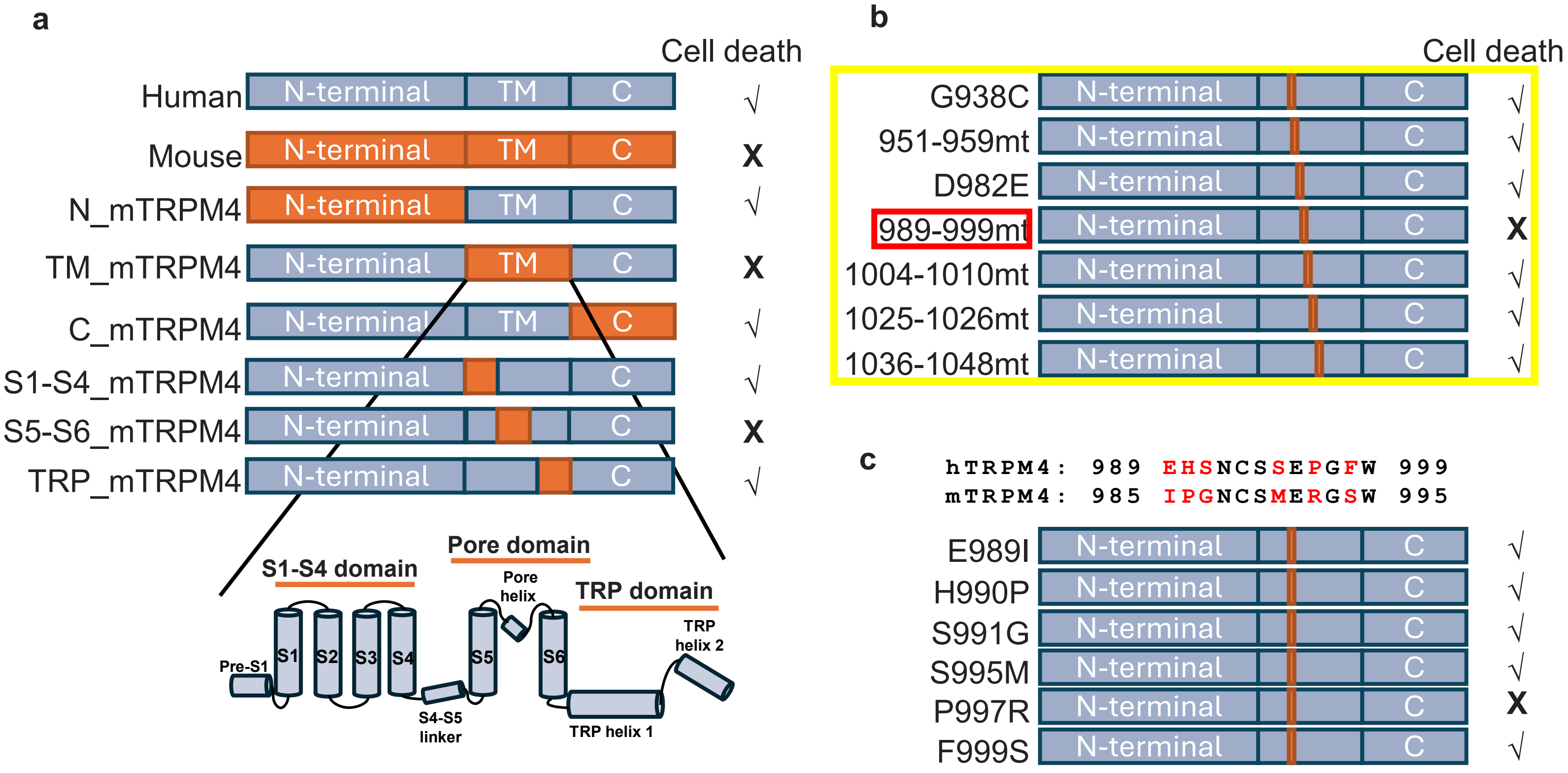


**Supplementary Fig. 6: Identification of a potential NC1-insensitive loss-of-function mutation in hTRPM4.** **a** Chimeras were generated on the background of hTRPM4 to identify the region important for the channel's sensitivity to NC1 activation. When the TM region (Pre-S1-TRP helices) or the pore domain (S5-S6) of hTRPM4 was replaced by the corresponding region from mTRPM4, the HeLa cells expressing the chimeras became resistant to NC1-induced necrosis. **b** Following this observation, a set of more refined chimera and single-mutation constructs within the pore domain of hTRPM4 was generated. A 11-residue region (residues 989-999) was shown to be important for hTRPM4 sensitivity to NC1, as its replacement with the equivalent residues from mTRPM4 (residues 985-995) compromises the mutant sensitivity to NC1-induced necrosis. **c** Further tests using single mutations (highlighted in red in the alignment) within this 11-residue region suggest that P997 of hTRPM4 (R993 in mTRPM4) may be important for the NC1 activation of hTRPM4.

**a**

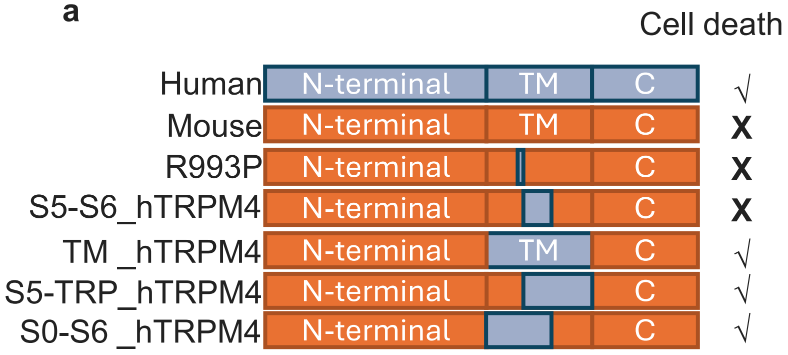


**b**

| Mutation | | Domain | Cell death |
|---|---|---|---|
| mTRPM4 R993P plus | S857A+T859S | S2-S3 linker | - |
| | L866A | S3 | - |
| | S1044G+H1047Q | S6 | - |
| | S1060R | TRP | + |
| | I897V+L900V | S4 | - |
| | I1021V+V1022I | S6 | - |
| | L1032V | S6 | - |
| | C934G | S5 | - |
| | F884Y+D885H | S3 | - |
| | H853R | S2-S3 linker | - |
| | R849S+H850Q | S2-S3 linker | - |
| | R841P+D844G | S2-S3 linker | - |
| | T804A+K805P | S1-S2 | - |
| | A778I+L780M | S0-S1 | - |
| | S769F+D770H | S0 | - |
| | R845H+P847S | S2-S3 linker | - |
| | G830S+W833G | S2 | - |
| | SVS807/8/9GSL | S1-S2 | - |
| | A794S+H795R | S1 | - |
| | L893I | S4 | - |
| | S765L+K766R | S0 | - |

**c**

| Mutation | | Domain | Cell death |
|---|---|---|---|
| mTRPM4 R993P/S1060R pus | S857A+T859S | S2-S3 linker | + |
| | L866A | S3 | + |
| | I897V+L900V | S4 | +++ |
| | F884Y+D885H | S3 | + |
| | H853R | S2-S3 linker | + |
| | R849S+H850Q | S2-S3 linker | + |
| | R841P+D844G | S2-S3 linker | + |
| | T804A+K805P | S1-S2 | + |
| | A778I+L780M | S0-S1 | + |
| | S769F+D770 | S0 | + |
| | R845H+P847S | S2-S3 linker | + |
| | G830S+W833G | S2 | + |
| | SVS807/8/9GSL | S1-S2 | + |
| | A794S+H795R | S1 | + |
| | L893I | S4 | + |
| | S765L+K766R | S0 | + |
| | L900V | S4 | +++ |

**Supplementary Fig. 7: Identification of potential NC1-sensitive gain-of-function mutations in mTRPM4. a** A second set of mutations was generated on the background of mTRPM4 to identify the gain-of-function mutations that would convey NC1 sensitivity to the mouse TRPM4. Interestingly, while P997 was shown to be important for NC1 sensitivity in hTRPM4, mutating the equivalent R993 to Pro in mTRPM4 is insufficient to convey the mutant-expressing HeLa cells sensitive to NC1-induced necrosis. We therefore generated another set of chimeras in which various parts of the mTRPM4 TM regions were replaced with the hTRPM4 counterparts.  Among those tested, swapping the S0-TRP (TM), S0-S6, or the S5-TRP region of mTRPM4 with that of hTRPM4 can render the channel sensitive to NC1, suggesting that some residues within S0-S4 and the TRP domain are important for NC1 specificity. **b** We then generated a set of double or triple mutations on the background of R993P and identified a second gain-of-function mutation, S1060R. **c** A further test with triple or quadruple mutations on the background of R993P/S1060R led to the identification of the third potential gain-of-function mutation, L900V.

For data shown in **b** and **c**, HeLa cells transiently transfected with different GFP-tagged mTRPM4 mutants were treated with 1 μM NC1 for 6 hours instead of 24 hours to distinguish the NC1 sensitivity among these mutants. The morphological changes of cells with GFP fluorescence were monitored using microscopy. “-”, no morphological change was observed in transfected cells; “+”, balloon-like morphology was observed among some of the transfected cells; “+++”, all transfected cells had balloon-like morphology. More than 100 cells with GFP fluorescence were analyzed in each group, and the assays were repeated in three independent experiments with similar results.

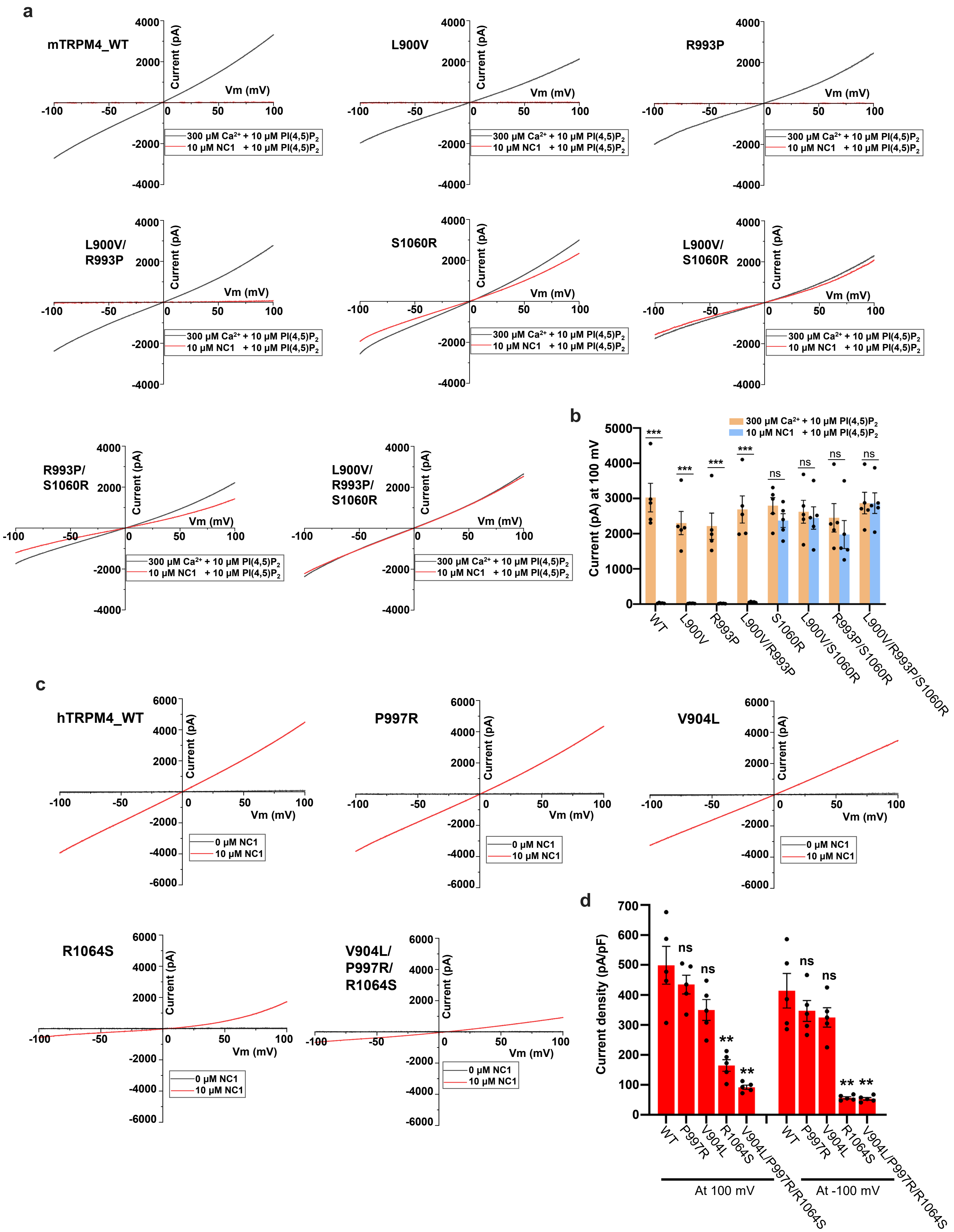

a
mTRPM4_WT
L900V
R993P
L900V/
R993P
S1060R
L900V/
S1060R
R993P/
S1060R
L900V/
R993P/
S1060R
Current (pA)
Vm (mV)
300 μM Ca2+ + 10 μM PI(4,5)P2
10 μM NC1 + 10 μM PI(4,5)P2
b
Current (pA) at 100 mV
***
ns
WT
L900V
R993P
L900V/R993P
S1060R
L900V/S1060R
R993P/S1060R
L900V/R993P/S1060R
c
hTRPM4_WT
P997R
V904L
R1064S
V904L/
P997R/
R1064S
0 μM NC1
10 μM NC1
d
Current density (pA/pF)
ns
**
WT
P997R
V904L
R1064S
V904L/P997R/R1064S
At 100 mV
At -100 mV

**Supplementary Fig. 8: Electrophysiology of mutations at the residues important for NC1 specificity. a** Sample I-V curves of wild-type mTRPM4 and its mutants at the three key residues. Currents were recorded in inside-out patches with the presence of 10 μM PI(4,5)$P_2$ diC8 and either 300 μM $Ca^{2+}$ or 10 μM NC1 in the bath (cytosolic). **b** Outward currents of wild-type mTRPM4 and its mutants recorded at +100 mV in inside-out patches with the presence of 10 μM PI(4,5)$P_2$ diC8 and either 300 μM $Ca^{2+}$ or 10 μM NC1 in the bath (cytosolic). Bars represent mean ± SEM of n=5 independent replicates (shown as dots). p-values were calculated using a two-sided Student's t-test and are provided in the Source Data file. *** represents $p < 0.001$. **c** Sample I-V curves of wild-type hTRPM4 and its single and triple mutants at the three key residues. Currents were recorded in whole-cell patches with 10 μM NC1 in the bath (extracellular). **d** Outward and inward current densities of the wild-type hTRPM4 and its mutants recorded at ±100 mV in whole-cell patches with 10 μM NC1 in the bath (extracellular). Bars represent mean ± SEM of n=5 independent replicates (shown as dots). p-values were calculated using a two-sided Student's t-test and are provided in the Source Data file. ** represents $p < 0.01$.

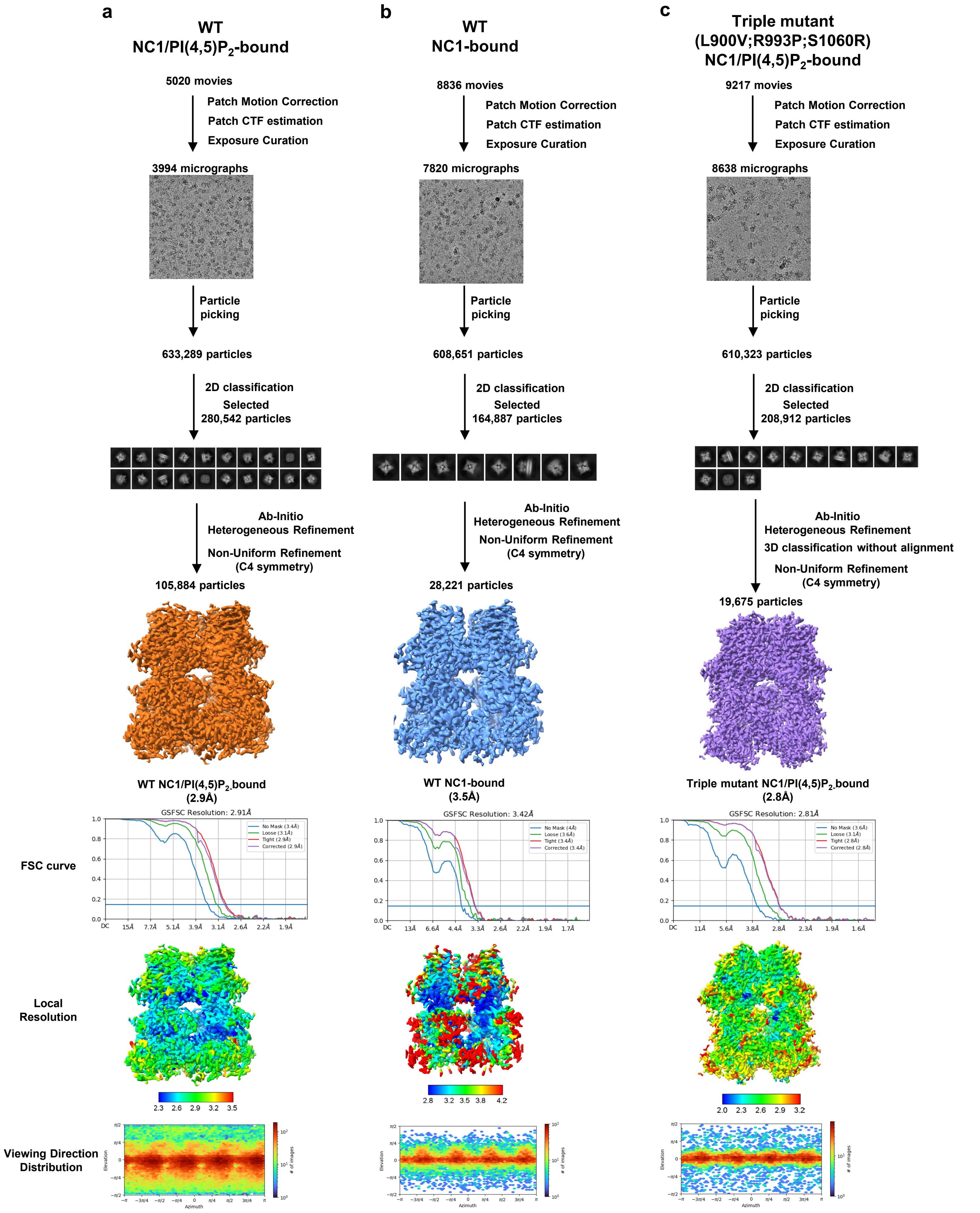


**Supplementary Fig. 9: Cryo-EM data processing scheme for the NC1-bound mTRPM4 samples.**

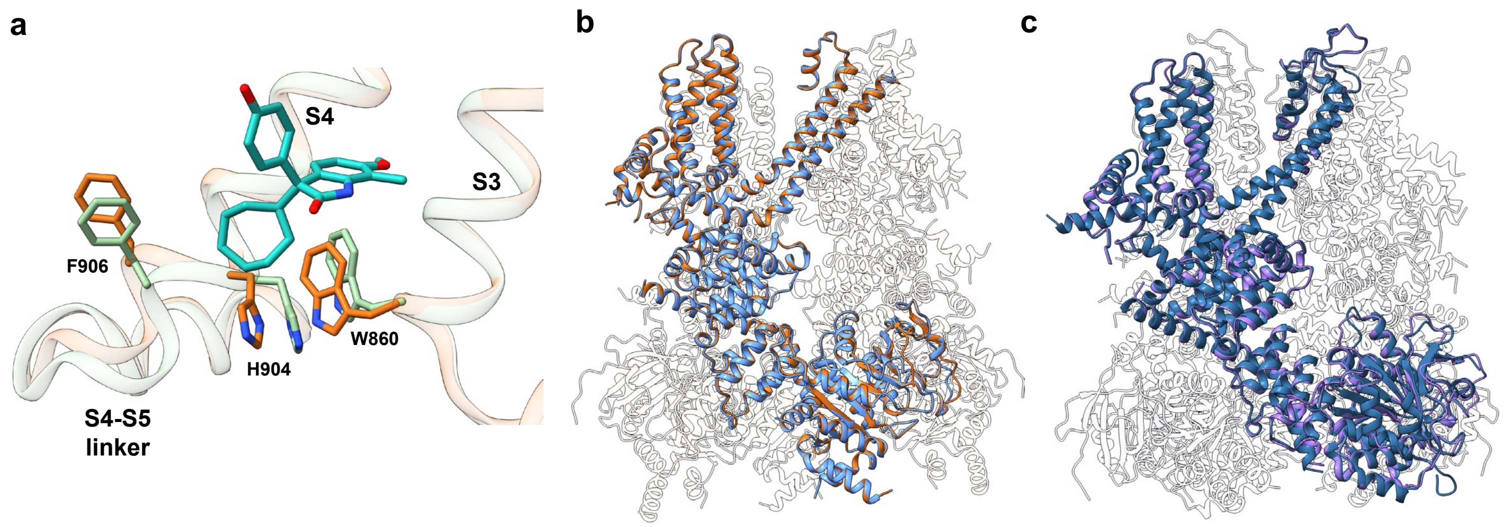


**Supplementary Fig. 10: Comparison of mTRPM4 structures with bound NC1. a** Local conformational changes within the S1-S4 domain between the NC1/PI(4,5)$P_2$-bound (light brown) and apo closed (light green) wild-type mTRPM4 structures. **b** Superposition of the NC1/PI(4,5)$P_2$-bound and NC1-bound wild-type mTRPM4 structures with the front subunits highlighted in brown and blue, respectively. The two structures are virtually identical. **c** Superposition of the NC1/PI(4,5)$P_2$-bound mTRPM4 triple mutant (L900V/R993P/S1060R) and NC1/PI(4,5)$P_2$-bound wild-type hTRPM4 structures with the front subunits highlighted in purple and navy blue, respectively. The two structures are virtually identical and represent an open conformation.

| | NC1 +EGTA | NC1/PI(4,5)$P_2$ | NC1/PI(4,5)$P_2$ +EGTA |
|---|---|---|---|
| | EMD-73526 PDB-9YVK | EMD-73527 PDB-9YVL | EMD-73528 PDB-9YVM |
| **Data collection and processing** | | | |
| Magnification | 165,000 | 105,000 | 81,000 |
| Voltage (kV) | 300 | 300 | 300 |
| Electron exposure ($e^-$/Å$^2$) | 60 | 60 | 60 |
| Defocus range (µm) | -0.9 - -2.2 | -0.9 - -2.2 | -0.9 - -2.2 |
| Pixel size (Å) | 0.738 | 0.834 | 0.857 |
| Symmetry imposed | C4 | C4 | C4 |
| Initial particle images (no.) | 1,285,163 | 891,246 | 2,294,749 |
| Final particle images (no.) | 112,765 | 32,516 | 79,688 |
| Map resolution (Å) | 2.69 | 2.78 | 2.73 |
| FSC threshold: 0.143 | | | |
| **Refinement** | | | |
| Model resolution (Å) | 2.66 | 2.75 | 2.71 |
| FSC threshold: 0.143 | | | |
| Map sharpening B factor (Å$^2$) | -80.5 | -85.1 | -88.6 |
| Model composition | | | |
| Non-hydrogen atoms | 30856 | 30712 | 30708 |
| Protein residues | 3860 | 3804 | 3804 |
| Ligands | NC1:4 | $Ca^{2+}$:4 PI(4,5)$P_2$:4 NC1:4 | PI(4,5)$P_2$:4 NC1:4 |
| B factors (Å$^2$) | | | |
| Protein | 31.25 | 92.23 | 74.82 |
| Ligand | 30.00 | 57.24 | 39.88 |
| R.m.s. deviations | | | |
| Bond lengths (Å) | 0.004 | 0.004 | 0.005 |
| Bond angles (°) | 0.557 | 0.535 | 0.661 |
| Validation | | | |
| MolProbity score | 0.99 | 1.12 | 1.14 |
| Clashscore | 2.13 | 3.24 | 3.52 |
| Poor rotamers (%) | 0 | 1.00 | 0 |
| Ramachandran plot | | | |
| Favored (%) | 98.39 | 98.19 | 98.49 |
| Allowed (%) | 1.61 | 1.81 | 1.51 |
| Disallowed (%) | 0 | 0 | 0 |

**Supplementary Table 1: hTRPM4 cryo-EM data collection, refinement and validation statistics**

| | WT<br>NC1/PI(4,5)$P_2$<br><br>EMD-73530<br>PDB-9YVO | WT<br>NC1<br><br>EMD-73529<br>PDB-9YVN | Triple mutant<br>NC1/PI(4,5)$P_2$<br><br>EMD-73531<br>PDB-9YVP |
|---|---|---|---|
| **Data collection and processing** | | | |
| Magnification | 81,000 | 165,000 | 165,000 |
| Voltage (kV) | 300 | 300 | 300 |
| Electron exposure ($e^-$/Å$^2$) | 60 | 60 | 60 |
| Defocus range (μm) | -0.9 - -2.2 | -0.9 - -2.2 | -0.9 - -2.2 |
| Pixel size (Å) | 0.857 | 0.735 | 0.735 |
| Symmetry imposed | C4 | C4 | C4 |
| Initial particle images (no.) | 633,289 | 608,651 | 610,323 |
| Final particle images (no.) | 105,884 | 28,221 | 19,675 |
| Map resolution (Å)<br>FSC threshold: 0.143 | 2.91 | 3.42 | 2.81 |
| **Refinement** | | | |
| Model resolution (Å)<br>FSC threshold: 0.143 | 2.88 | 3.39 | 2.78 |
| Map sharpening B factor (Å$^2$) | -101.1 | -88.5 | -60.0 |
| Model composition | | | |
| Non-hydrogen atoms | 27924 | 27636 | 30036 |
| Protein residues | 3464 | 3456 | 3732 |
| Ligands | $Ca^{2+}$:8<br>PI(4,5)$P_2$:4<br>NC1:4 | NC1:4 | $Ca^{2+}$:8<br>PI(4,5)$P_2$:4<br>NC1:4 |
| B factors (Å$^2$) | | | |
| Protein | 82.71 | 103.37 | 58.65 |
| Ligand | 58.50 | 58.86 | 57.62 |
| R.m.s. deviations | | | |
| Bond lengths (Å) | 0.005 | 0.006 | 0.003 |
| Bond angles (°) | 0.666 | 0.733 | 0.517 |
| Validation | | | |
| MolProbity score | 1.21 | 1.08 | 1.02 |
| Clashscore | 3.62 | 2.90 | 2.39 |
| Poor rotamers (%) | 0.68 | 0 | 0 |
| Ramachandran plot | | | |
| Favored (%) | 97.76 | 98.73 | 98.26 |
| Allowed (%) | 2.24 | 1.27 | 1.74 |
| Disallowed (%) | 0 | 0 | 0 |

**Supplementary Table 2: mTRPM4 cryo-EM data collection, refinement and validation statistics**
